# Motor cortical signals predict finger velocity beyond task geometry

**DOI:** 10.64898/2026.08.04.742876

**Authors:** Zonghan Du, Zhong Yuan Lai, Liang Hu, Tangwei Ye

## Abstract

Finger movements toward the same target can differ in speed. How much does motor cortical activity add to a velocity prediction once the decoder knows the starting position and the intended target? We compared predictions of finger velocity on held-out trials with and without spiking-band power from motor cortex. In 18 sessions from one macaque, neural activity reduced mean squared error by 4.92% relative to a baseline that used the target, starting position and time within the trial. The reduction was 4.63% in 20 additional sessions from the same animal, with improvement in every session. Giving the baseline the preceding 100 ms of measured finger position reduced the neural gain in every session. Mean error reductions fell to 2.75% and 2.05%, respectively. In all 20 additional sessions, neural activity from the same trial gave more accurate predictions than activity substituted from a closely matched trial. In simulations, an incomplete model of task geometry could also produce this pairing effect. Motor cortical activity predicts variation in finger velocity beyond the movement expected from task geometry, and part of that variation is also predictable from recent finger position.

## Introduction

Two movements can have the same starting position and target yet differ in when the fingers accelerate or slow down. Macaques have used motor cortical activity to control two virtual finger groups independently in real time [1]. In the Long-term Intracortical Neural Activity and Kinematics (LINK) dataset, Temmar and colleagues decoded the velocity of these finger groups across recording days [2]. A decoder can predict the movement typical of a target and the deviations specific to a trial. Its overall accuracy combines these contributions. For a decoder that already receives the target, the relevant quantity is the change in prediction error when neural activity is added.

In task-performing mice, instructed and uninstructed movements explained much of the single-trial cortical activity [3]. In spontaneously behaving mice, recorded movements predicted widespread activity, including activity in visual cortex [4]. Truccolo and colleagues separated the contributions of spike history, ensemble activity and external covariates to spiking [5]. De Heer and colleagues used variance partitioning to estimate the unique and shared contributions of speech features to cortical responses [6]. Here velocity is the predicted quantity, and the added predictors are neural measurements. A movement target can guide a trajectory decoder [7], and an autoregressive decoder can use earlier kinematics to predict the next movement [8]. These two sources of information provide separate reference points for measuring what a cortical recording contributes.

We used repeated finger movements in LINK to ask how much spiking-band power adds at each reference point. Spiking-band power summarizes local spiking activity on an electrode and has been used to decode finger movement [9]. If recent position and cortical activity track some of the same deviations from the typical movement, recent position should reduce the extra improvement from cortical activity. Spiking-band power improved predictions beyond task geometry in each of 38 sessions from one macaque. Across the two sets of sessions, recent measured finger position reduced the mean improvement from about 5% of baseline error to about 2–3%. We then used trial replacement to test whether prediction depended on which trial supplied the neural activity. Continuous reaching in two other macaques provided a separate test of how recent position and recording time affected that comparison.

## Results

### Motor cortical activity improves predictions beyond task geometry

The macaque controlled its index finger and its three grouped fingers (middle, ring and small) in center-out and random-target movements. We reserved whole trials for testing and fitted the models on the other trials. For each held-out trial we predicted both components of finger velocity in 20 ms bins. The *geometry baseline* used the target, starting position and time within the trial to predict the movement typical of that geometry. We gave the model 160 ms of preceding motor cortical spiking-band power to predict the remaining velocity error. Figure 1a shows the resulting change in the predicted velocity trace for one held-out trial.

**Figure 1.**
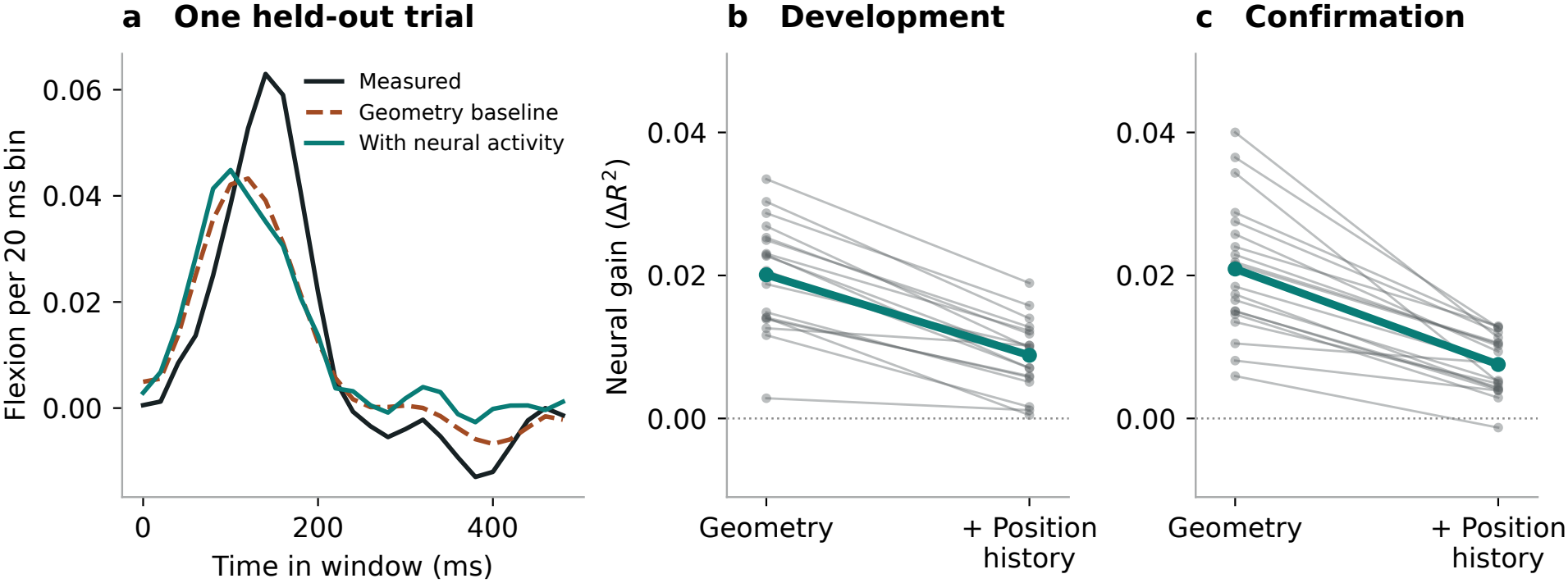
Motor cortical spiking-band power improves finger velocity predictions. **a**, Measured and predicted index-finger velocity in a held-out random-target trial on 21 September 2022. The session and trial were selected by median prediction improvement (Methods). The two predictions use the same geometry baseline. **b**,**c**, Gain from adding neural activity before and after giving the baseline recent finger position. Each line connects the two estimates for one session. The heavy line connects session means. Zero marks equal error with and without neural activity.

In the 18 development sessions, adding neural activity reduced mean squared error by 4.92% relative to the geometry baseline. Mean root mean squared error fell from 0.01407 to 0.01372 flexion units per 20 ms bin. The improvement was positive in every session (Fig. 1b). Measured as a fraction of the variance in held-out velocity, the mean gain was Δ*R*^2^ = 0.02009 (95% interval across recording dates, 0.01586 to 0.02404). Thus Δ*R*^2^ = 0.02009 corresponds to about two percentage points of held-out velocity variance. The 4.92% figure uses the baseline’s remaining error as its denominator. In the session illustrated in Fig. 1a, mean squared error fell from 0.000380 to 0.000360 in squared flexion units per bin. That change corresponds to Δ*R*^2^ = 0.02067 and a 5.20% reduction in baseline error.

Twenty additional sessions from the same macaque provided a confirmation set. Session selection used recording metadata before scores were available. Spiking-band power reduced mean squared error relative to the geometry baseline by 4.63% on average and improved prediction in all 20 sessions (mean Δ*R*^2^ = 0.02090).

### Recent finger position reduces the contribution of neural activity

The *position-history baseline* also received the previous five measurements of finger position, spanning 100 ms. It could therefore distinguish movements headed for the same target that had already taken different trajectories. Adding spiking-band power to this baseline reduced mean squared error by 2.75% in development and 2.05% in confirmation. Within each of the 38 sessions, recent finger position reduced the gain from adding spiking-band power relative to the geometry baseline (Fig. 1b,c). The corresponding mean Δ*R*^2^ was 0.00883 (95% interval 0.00628 to 0.01097) and 0.00755. Prediction improved in 18 of 18 and 19 of 20 sessions, respectively (Fig. 1b,c).

Threshold-crossing rate from the same electrodes gave a second measure of neural activity in the development sessions. It reduced mean squared error by 2.75% relative to the geometry baseline and by 1.37% relative to the position-history baseline. Each baseline gave a positive gain in 17 of 18 sessions. The ordering of the two baselines therefore held for both neural signals. Time within the trial alone explained less than 1% of velocity variance in either task. Adding geometry raised baseline *R*^2^ to 0.570 in center-out and 0.600 in random-target sessions. The ordering of center-out and random-target sessions by neural gain depended on the baseline (Supplementary Tables S3 and S4).

### Same-trial neural activity predicts finger velocity more accurately

If the correction to a velocity prediction depends on activity specific to that trial, substituting activity from another trial should change the prediction. For each held-out trial we kept its velocity and baseline prediction fixed. We replaced its neural residual with the residual from another held-out trial with the same movement direction and the closest baseline inputs. A neural residual is the recorded activity minus the activity predicted from that trial’s own baseline inputs.

With the geometry baseline, replacement changed mean Δ*R*^2^ from 0.02009 to −0.00714 in development and from 0.02090 to −0.00762 in confirmation. With the position-history baseline, replacement gave mean gains of −0.00384 and −0.00498, respectively. For both baselines the same trial gave the better prediction in every confirmation session (Fig. 2a). Replacement itself improved on the baseline in three of 20 confirmation sessions for each baseline. Two simpler baselines also gave positive replacement gains. The mean was 0.10498 for time alone in random-target sessions and 0.01294 for geometry without time interactions in center-out sessions (Supplementary Table S3). Thus a substituted trial can improve on the baseline even when activity from the original trial gives the better prediction.

**Figure 2.**
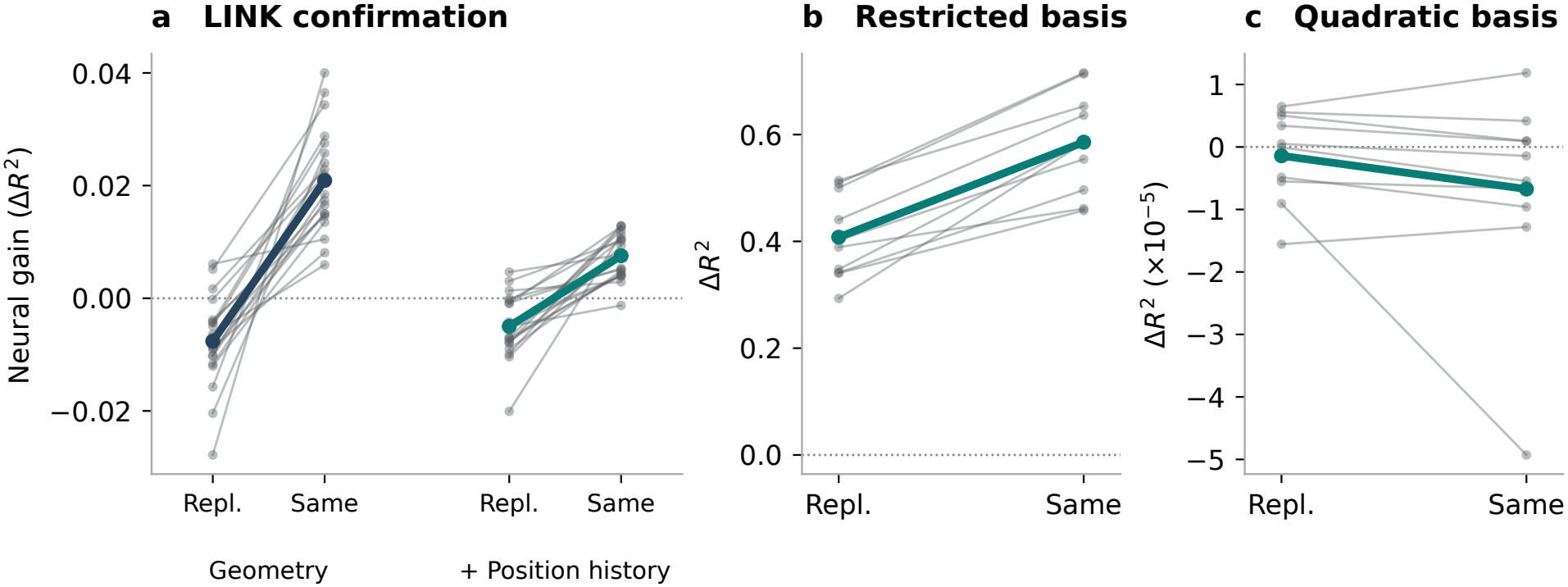
Trial replacement tests which neural recording supplies the correction. **a**, Each line joins the gain with substituted and same-trial neural activity in one confirmation session, shown separately for the two baselines. Repl. denotes the matched replacement trial and Same denotes the target trial. **b**,**c**, In the simulation, neural activity and velocity have independent noise conditional on geometry and time. Each line joins the replacement and same-trial gains for one seed. The quadratic baseline in **c** includes the terms used to generate the data. Its vertical scale resolves changes of 10^−5^ in Δ*R*^2^. Heavy lines show means.

To establish what this comparison can identify, we generated movements and neural activity that depended on the same observed geometry and time, with independent noise. Across ten simulations a baseline missing quadratic geometry terms gave a mean same-trial gain of 0.58571 and a mean replacement gain of 0.40784. The same trial gave the better prediction in all ten. Including the quadratic terms, computed from the same inputs, brought mean same-trial gain to −0.00000673 (Fig. 2b,c). A dependence on trial pairing can therefore arise when neural activity supplies a feature of geometry that the baseline fails to fit.

### Recent cursor position and prediction accuracy in two reaching macaques

We applied the same comparison to continuous reaching in two other macaques, Indy and Loco [10]. Each animal contributed ten dates. The task baseline used the current target, previous target and time since the target changed. The position-history baseline also used 100 ms of measured cursor position. We trained and evaluated the models separately within each recording, using earlier blocks for training and later blocks for testing.

The primary quadratic models gave mean task and position-history gains of 0.2737 and 0.0572 in Indy. In Loco the corresponding gains were 11.7338 and 0.0276. One Loco session accounted for most of the large task gain. The target remained unchanged for up to 48.896 s during testing, compared with 8.812 s during training. On that date the quadratic task baseline and the full prediction had *R*^2^ values of −330.08 and −214.00. Their difference of 116.07 reflects two inaccurate predictions. Capping elapsed time at its training maximum in a post hoc comparison reduced the full prediction’s root mean squared error from 497.2 to 45.6 mm/s.

The prespecified linear models tested the same inputs without quadratic terms. Recent cursor position reduced the gain from adding motor cortical activity on all 20 dates. Mean Δ*R*^2^ fell from 0.2695 to 0.0612 in Indy and from 0.1830 to 0.0180 in Loco (Fig. 3a).

**Figure 3.**
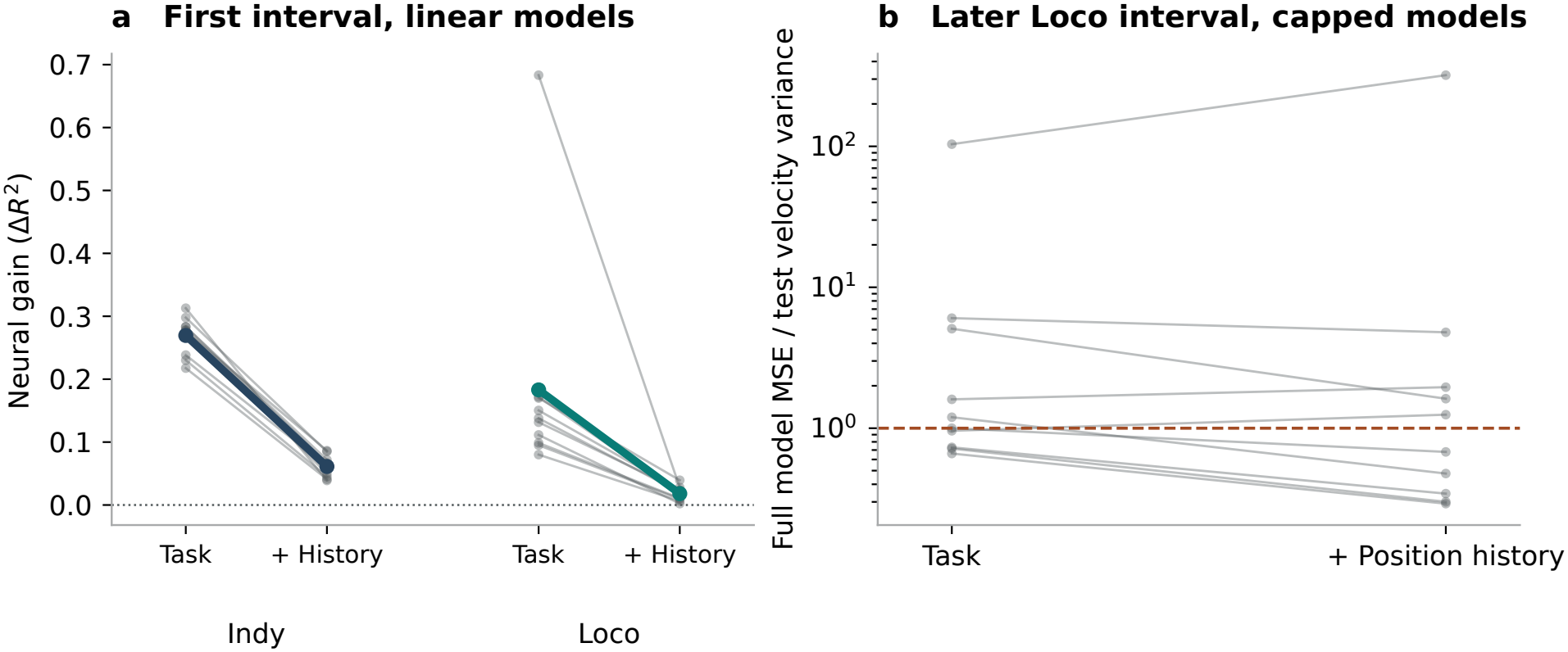
The change in error depends on recent cursor position and on when the model is tested. **a**, Each line joins linear-baseline gains for one date in the first recording interval. **b**, Full prediction error on later Loco blocks divided by the error of the test-interval mean, using the capped quadratic models. Values above one correspond to negative *R*^2^. Each line is one recording date. Supplementary Tables S22–S29 give all model forms and dates.

A later interval in the same ten Loco recordings tested how well fixed predictions held up over time. In the interval from 602 to 1,100 s, the capped quadratic full prediction had negative *R*^2^ on six dates with task inputs and five with position history (Fig. 3b). Negative *R*^2^ means that the prediction had greater squared error than the mean velocity of that test interval. Same-block activity still outperformed replacement in all 20 capped comparisons. For each of the three baseline forms, the interval across dates for the difference in neural gain between task inputs and position history included zero. On these later blocks, the paired neural recording predicted velocity more accurately than its replacement even on dates with large absolute errors.

## Discussion

In the LINK finger task, adding motor cortical spiking-band power removed about 5% of the error left by task geometry. When the decoder also knew where the fingers had recently been, spiking-band power removed about 2–3% of the remaining error. Recent position can distinguish two movements headed for the same target when one has progressed further than the other. The smaller neural gain in every session is consistent with cortical activity and recent position tracking some of the same trial-to-trial variation.

In macaque reaching, motor cortical population dynamics have improved single-trial decoding [11], and the dimensions along which population activity rotates vary across reaches [12]. Motor cortical neurons also respond to rapid sensory feedback following perturbations of the arm [13]. Motor commands and sensory feedback could therefore both contribute to the finger velocity correction. The simulation gives a second explanation for a pairing effect. Even when geometry drives every shared fluctuation, a restricted baseline can leave predictable structure in the neural residual.

A decoder with an external position sensor can use recent measured position directly. A decoder that feeds back its own predicted cursor position carries the error in that estimate into its next prediction. The LINK percentages quantify the contribution of neural activity when position is measured. In continuous reaching, the full model on one Loco date reduced the squared error of the task baseline yet still missed the measured velocity by 497.2 mm/s in root mean squared error. Reporting both errors in movement units alongside the neural gain reveals the accuracy of the prediction an interface would use.

Across 38 finger-movement sessions, knowing the target and starting position left a reproducible contribution from same-trial motor cortical activity. Knowing the recent finger trajectory reduced the neural gain in every session.

## Methods

### LINK recordings and velocity targets

LINK contains 312 recording sessions from a macaque with arrays implanted in right precentral gyrus [2, 14]. The development set comprised all nine dates with both a center-out and a random-target session. For confirmation we excluded those sessions, sorted the remaining recordings within task by date and asset identifier, and selected ten evenly spaced interior ranks per task. Each session contributed 375 usable trials. We kept each trial intact and stratified the 70/30 training/test split by the direction from start to target. The resulting sets contained 262 training trials and 113 test trials per session.

After the history window, each trial provided 25 predictions over 500 ms. LINK reports the positions of two finger groups in flexion units and velocity in flexion units per 20 ms bin. The source dataset describes averaging 1 kHz position measurements in non-overlapping 20 ms bins before differencing [2]. In all 38 files, both released velocity components at each checked interior bin *k* equalled *p*_*k*+1_ − *p*_*k*_. Neural inputs were the 96-channel spiking-band power in bins *k* − 8 through *k* − 1. The position-history baseline also used both positions in bins *k* − 5 through *k* − 1. The inputs end before the two position bins defining the velocity target. The release specifies bin order but gives no within-bin timestamp convention or hardware latency.

For each time bin we represented normalized time within the analysis window by a constant and four sine/cosine pairs. The geometry baseline included this time basis, its interactions with eight movement-direction categories, and target position, starting position and displacement from start to target, each with its time interactions. This gave 141 features. The position-history baseline appended ten position values, giving 151 features. The threshold-crossing analysis substituted threshold-crossing rates on the same 96 channels for spiking-band power.

### Estimating the improvement from neural activity

For baseline inputs *C*, velocity *Y* and neural measurements *X*, we fitted separate ridge regressions *m*_*Y*_ (*C*) and *m*_*X*_(*C*). The first predicts velocity from the baseline inputs. The second predicts the neural activity expected from those same inputs. A third ridge regression *h* predicted the remaining velocity error from the remaining neural activity. The full prediction was

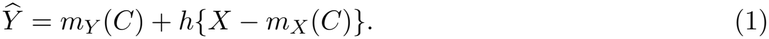

Leave-one-covariate-out prediction measures the change in test error after omitting a predictor and refitting [15]. Here the added predictor block is the preceding neural activity across all electrodes. We estimated the baseline-predicted component of each neural input from training data, following cross-validated approaches to adjustment for measured covariates [16, 17]. We trained *h* on out-of-fold residuals from five folds of whole training trials. Each nuisance regression selected its penalty on an inner 80/20 split and refitted on the corresponding training subset. A separate three-fold loop selected the penalty for *h* by repeating the nuisance fits and their penalty selection inside each training fold [18]. Each scaler used the training data for its own fit. All regressions considered penalties 0.1, 1, 10, 100, 10^3^, 10^4^, 10^5^. The selected model used the full outer training set for its final nuisance fits. Supplementary Methods give the fold construction and timing for each dataset.

We averaged squared error over the two velocity components and all held-out time bins within a session. Let *E*_*B*_ and *E*_*F*_ be the resulting baseline and full mean squared errors, and let *V* be the average of the two test-component variances. For *n* held-out time bins with two velocity components, the mean squared error of a prediction is

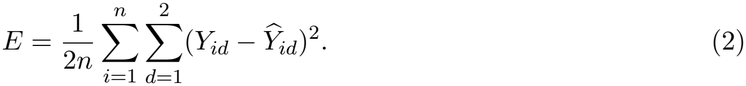

The percentage of baseline error removed uses *E*_*B*_ as the denominator. Neural gain uses *V* as the denominator.

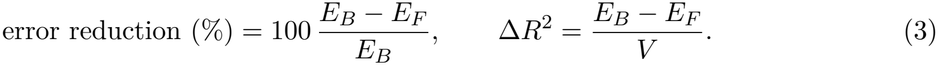

We computed both quantities within each session and then averaged sessions with equal weight. Development intervals resampled the nine recording dates, keeping the two tasks together. Confirmation counts refer to individual sessions from the same animal. To illustrate the LINK predictions we used the session at the lower median geometry-baseline gain and the trial at the lower median improvement in mean squared error within that session.

### Trial replacement and simulations

We matched each held-out trial to a different held-out trial of the same movement direction. Among eligible trials we selected the closest complete trajectory of baseline inputs by mean squared distance after scaling with outer-training statistics. The lower trial index broke ties. We replaced *X* −*m*_*X*_(*C*) with the source trial’s neural residual while keeping the target trial’s baseline prediction and measured velocity fixed. Matching uses the full held-out trajectory and is an offline comparison.

For the counterexample in Fig. 2b,c, the simulation generated neural activity and velocity from functions of the same two continuous geometry variables and trial phase. Independent noise processes gave conditional independence by construction. The restricted baseline contained the 17 linear geometry and phase terms. The expanded baseline used all 35 generating terms, including quadratic geometry and its interactions with phase. Both fits used identical observations and splits within each of ten fixed seeds. A separate calibration injected a known shared trial deviation at three strengths (Supplementary Methods and Tables S17 and S21).

### Continuous reaching and evaluation at a later time

We selected ten dates for each macaque from the Indy and Loco reaching dataset [10]. We sorted dates, retained the first file per date, and took ten evenly spaced date indices rounded down. The source contained 34 eligible recording dates for Indy and ten for Loco. We used the first 600 s, or the entire file for shorter recordings of at least 300 s. We pooled sorted and unsorted spikes within each M1 channel. We averaged each set of five native 250 Hz cursor positions to form a 20 ms bin, then divided successive position differences by their timestamp differences to obtain velocity in mm/s. Neural and position histories spanned 100 ms and ended 80 ms before the target bin. Task inputs used the last available values of the current target, previous target and elapsed time since the target changed.

Non-overlapping 2 s blocks each contributed 90 predictions. Within a recording, the first 70% of physical blocks (rounded down) formed the training period. We skipped the next block and evaluated the remaining eligible blocks. The primary models included linear and quadratic terms in their inputs. The prespecified sensitivity used linear terms. After a large extrapolation error appeared, a post hoc comparison capped elapsed time at the maximum in each training subset before constructing quadratic terms. The later evaluation used a protocol fixed before scoring the new interval. All six model specifications retained their original training blocks and fitted predictions. We skipped 600–602 s and scored the next 249 blocks through 1,100 s on each Loco date. Within-session intervals resampled blocks. Across-date intervals resampled dates within animal. The 2 s blocks have disjoint measurements but can remain temporally correlated.

## Supporting information

Source data and analysis code

## Data and code availability

The LINK finger recordings [14], the MC_Maze and MC_RTT recordings from the Neural Latents Benchmark [19, 20, 21], and the Indy and Loco reaching recordings [10] are available under CC BY 4.0. The supplementary data and code include the session-level estimates, the data and scripts for each figure, the model-fitting code and the held-out predictions for all 346 linear fits in the finger, benchmark and calibration analyses. The accompanying guide links each analysis to its inputs and software.

## Ethics

We analysed public nonhuman-primate recordings collected by the original investigators. The LINK dataset lists animal-use approval PRO0001815.

## Acknowledgements

We thank the investigators who collected and shared the LINK, Neural Latents Benchmark, and Indy and Loco reaching datasets. OpenAI Codex assisted with code development, analysis checks, figure preparation and manuscript editing.

## Author contributions

Zonghan Du: conceptualization, methodology, software, formal analysis, investigation, visualization, and writing—original draft. Zhong Yuan Lai: methodology, formal analysis, supervision, and writing—review and editing. Liang Hu: conceptualization, methodology, investigation, supervision, and writing—review and editing. Tangwei Ye: software, formal analysis, validation, and writing—review and editing.

## References

[1] Nason SR, Mender MJ, Vaskov AK, et al. Real-time linear prediction of simultaneous and independent movements of two finger groups using an intracortical brain-machine interface.Neuron. 2021;109:3164–3177.e8. 10.1016/j.neuron.2021.08.009

[2] Temmar H, Wang Y, Gill N, et al. Long-term Intracortical Neural activity and Kinematics (LINK): an intracortical neural dataset for chronic brain-machine interfaces, neuroscience, and machine learning. NeurIPS Datasets and Benchmarks. 2025. https://papers.nips.cc/paper_files/paper/2025/hash/cb463f73a35802996546ac8e8b1b2743-Abstract-Datasets_and_Benchmarks_Track.html

[3] Musall S, Kaufman MT, Juavinett AL, Gluf S, Churchland AK. Single-trial neural dynamics are dominated by richly varied movements. Nature Neuroscience. 2019;22:1677–1686. 10.1038/s41593-019-0502-4

[4] Stringer C, Pachitariu M, Steinmetz N, Reddy CB, Carandini M, Harris KD. Spontaneous behaviors drive multidimensional, brainwide activity. Science. 2019;364:eaav7893. 10.1126/science.aav7893

[5] Truccolo W, Eden UT, Fellows MR, Donoghue JP, Brown EN. A point process framework for relating neural spiking activity to spiking history, neural ensemble, and extrinsic covariate effects. Journal of Neurophysiology. 2005;93:1074–1089. 10.1152/jn.00697.2004

[6] de Heer WA, Huth AG, Griffiths TL, Gallant JL, Theunissen FE. The hierarchical cortical organization of human speech processing. Journal of Neuroscience. 2017;37:6539–6557. 10.1523/JNEUROSCI.3267-16.2017

[7] Shanechi MM, Williams ZM, Wornell GW, Hu RC, Powers M, Brown EN. A real-time brain-machine interface combining motor target and trajectory intent using an optimal feedback control design. PLoS ONE. 2013;8:e59049. 10.1371/journal.pone.0059049

[8] Fisher J, Black M. Motor cortical decoding using an autoregressive moving average model. Proceedings of the IEEE EMBS. 2005:2130–2133. 10.1109/IEMBS.2005.1616881

[9] Nason SR, Vaskov AK, Willsey MS, et al. A low-power band of neuronal spiking activity dominated by local single units improves the performance of brain–machine interfaces. Nature Biomedical Engineering. 2020;4:973–983. 10.1038/s41551-020-0591-0

[10] O’Doherty JE, Cardoso MMB, Makin JG, Sabes PN. Nonhuman Primate Reaching with Multichannel Sensorimotor Cortex Electrophysiology. Zenodo; 2020. Fixed record 3854034. 10.5281/zenodo.3854034

[11] Kao JC, Nuyujukian P, Ryu SI, Churchland MM, Cunningham JP, Shenoy KV. Single-trial dynamics of motor cortex and their applications to brain–machine interfaces. Nature Communications. 2015;6:7759. 10.1038/ncomms8759

[12] Sabatini DA, Kaufman MT. Reach-dependent reorientation of rotational dynamics in motor cortex. Nature Communications. 2024;15:7007. 10.1038/s41467-024-51308-7

[13] Pruszynski JA, Kurtzer I, Nashed JY, Omrani M, Brouwer B, Scott SH. Primary motor cortex underlies multi-joint integration for fast feedback control. Nature. 2011;478:387–390. 10.1038/nature10436

[14] Temmar H, Gill N, Wang Y, et al. LINK: Long-Term Intracortical Neural Activity and Kinematics. DANDI Archive; 2025. Version 0.251023.2336. 10.48324/dandi.001201/0.251023.2336

[15] Lei J, G’Sell M, Rinaldo A, Tibshirani RJ, Wasserman L. Distribution-free predictive inference for regression. Journal of the American Statistical Association. 2018;113:1094–1111. 10.1080/01621459.2017.1307116

[16] Snoek L, Miletić S, Scholte HS. How to control for confounds in decoding analyses of neuroimaging data. NeuroImage. 2019;184:741–760. 10.1016/j.neuroimage.2018.09.074

[17] Chyzhyk D, Varoquaux G, Milham M, Thirion B. How to remove or control confounds in predictive models, with applications to brain biomarkers. GigaScience. 2022;11:giac014. 10.1093/gigascience/giac014

[18] Varma S, Simon R. Bias in error estimation when using cross-validation for model selection.BMC Bioinformatics. 2006;7:91. 10.1186/1471-2105-7-91

[19] Pei F, Ye J, Zoltowski DM, et al. Neural Latents Benchmark ‘21: evaluating latent variable models of neural population activity. Proceedings of the Neural Information Processing Systems Track on Datasets and Benchmarks. 2021;1. https://datasets-benchmarks-proceedings.neurips.cc/paper_files/paper/2021/hash/979d472a84804b9f647bc185a877a8b5-Abstract-round2.html

[20] Churchland M, Kaufman M. MC_Maze_Medium: macaque primary motor and dorsal premotor cortex spiking activity during delayed reaching. DANDI Archive; 2022. Version 0.220113.0408. 10.48324/dandi.000139/0.220113.0408

[21] O’Doherty J. MC_RTT: macaque motor cortex spiking activity during self-paced reaching. DANDI Archive; 2024. Version 0.241017.1444. 10.48324/dandi.000129/0.241017.1444

