## Supplementary methods and tables for "Motor cortical signals predict finger velocity beyond task geometry"

### What the gain measures

Let $Y$ be velocity, $X$ the neural measurements and $C$ the baseline inputs. Under squared error, the population improvement obtained by adding $X$ to an unrestricted predictor of $Y$ is

$$\mathbb{E}\big\|Y - \mathbb{E}[Y \mid C]\big\|^2 - \mathbb{E}\big\|Y - \mathbb{E}[Y \mid X, C]\big\|^2 = \mathbb{E}\big\|\mathbb{E}[Y \mid X, C] - \mathbb{E}[Y \mid C]\big\|^2. \tag{S1}$$

The squared Euclidean norm sums the squared errors of the two components. The expectation averages this loss over movements from the population. This identity follows by expanding the square and taking the conditional expectation of the cross term given $(X, C)$. It concerns the conditional mean of velocity. The gain from two fitted models can be negative on held-out data. Our $\Delta R^2$ divides their observed difference in mean squared error by the average test-component variance.

Adding baseline inputs can either increase or decrease the population gain. For instance, if $X$ and $Y$ are independently noisy measurements of the same underlying position, observing that position removes the predictive contribution of $X$. Conversely, let $X$ and $C$ be independent fair bits and $Y$ their exclusive-or. Observing $C$ makes $X$ predictive of $Y$, although $X$ alone leaves the mean of $Y$ unchanged. The decrease observed in the two LINK cohorts is therefore an empirical result.

### Recording windows and prediction models

LINK uses a normalized within-window time $\tau \in [0, 1)$ and the basis $1, \sin(r\pi\tau), \cos(r\pi\tau)$ for $r = 1, \ldots, 4$. Tables S1–S4 give the fixed settings, dimensions, and progression from time alone to the position-history baseline. The initial position comes from the trial start. Recorded finger position and velocity have matching timestamps with spiking-band power (SBP) and threshold-crossing rate (TCR). The history for velocity bin $k$ ends at $k - 1$. The target is the difference of position bins $k$ and $k + 1$. For the public LINK acquisition and binning description, see Appendix A.3 of the LINK dataset paper. Neither a within-bin timestamp convention nor a measured hardware latency is available from these checks.

Five folds of outer training trials produce the out-of-fold residuals for the final neural regression. Within each of three separate folds for choosing its ridge penalty, we repeat the five-fold construction on that fold's training trials. Two nuisance regressions fitted on those training trials transform the validation trials. We select the penalty with the best mean variance-weighted residual $R^2$ across the three folds, taking the smaller penalty on a tie. Every nuisance fit selects its own ridge penalty and estimates its own predictor means and scales. The inner holdout uses 20% of trials, enlarged as needed in small benchmark folds to represent each condition. Final nuisance regressions use all outer training trials. Date-level LINK intervals resample pairs of center-out and random-target sessions 10,000 times. Within-session intervals resample whole held-out trials 500 times within condition, with the predictions and source assignments fixed.

For the main estimates, we refitted the nuisance models within each fold used to select the neural regression penalty. A comparison used the same outer splits, inputs and seeds but computed nuisance residuals once before penalty selection. The two procedures selected different penalties in

28 of 460 fits. In the 20 confirmation sessions, the mean geometry-baseline gain was 0.02090 with fold-specific residuals and 0.02158 with reused residuals. The accompanying source data include held-out predictions for all 346 linear fits and scores and penalty-selection records for 114 nonlinear fits.

Tables S5 and S8–S14 report separate sensitivity analyses of baseline form, neural timing, train/test split and trial matching. In the linear residual regressions for these comparisons, one inner validation split selected the penalties. Six of nine sensitivity tables agreed with refits from the recordings to within $10^{-12}$ in the cells compared. In the other three, the largest difference among comparable cells was $1.64 \times 10^{-6}$. The original number of within-session bootstrap draws for the quadratic models is unknown, so 288 interval endpoints could not be reproduced exactly. Five additional random-feature diagnostics for a continuous-context simulation depend on a map seed that was not recorded.

### Matching, alternative predictors and benchmark recordings

For trial replacement, the target keeps its baseline prediction. Its source supplies neural measurements minus their expectation given the source's own inputs. Matching uses the squared distance between whole baseline trajectories, standardized against outer training data. One hundred within-condition permutations used randomly ordered trial indices and a cyclic shift without fixed points. The geometry baseline gave a negative mean permutation gain in each development session. Across-session mean permutation gain was $-0.0142$. In the five-baseline comparison, replacement gain was positive for the time-only baseline in random-target sessions (0.10498) and for additive geometry in center-out sessions (0.01294). Tables S3 and S14 give the individual comparisons and source ranks.

Two benchmark recordings illustrate the effect of different available inputs. MC_Maze-medium comprises 250 trials from one session in Jenkins. A template used 27 reach/barrier conditions and time from measured movement onset. Five neural bins over 152 units ended 80 ms before velocity. The release-defined split comprised 188 training and 62 validation trials. The template had $R^2 = 0.9053$. Adding neural activity reduced root mean squared error from 81.76 to 81.53 mm/s ($\Delta R^2 = 0.00051$). MC_RTT comprises 1,080 analysis blocks in one session in Indy. The release-defined split comprised 810 training and 270 validation blocks. Five neural bins over 130 units shared their end time with the position history. Adding neural activity to this baseline reduced root mean squared error from 39.78 to 36.19 mm/s ($\Delta R^2 = 0.06846$). The retrospective octant of mean target-minus-cursor displacement determined the strata and replacement pools in MC_RTT. The predictor itself used the specified lagged inputs. The two recordings have different tasks, animals, baseline inputs and event alignment. We used their released velocity series. Upstream differentiation and interpolation were not independently reconstructed.

Alternative models tested which parts of the LINK result depended on model form. They included joint ridge regression with separate penalties for baseline and neural features, quadratic geometry, and radial-basis approximations constructed with random Fourier features. The nonlinear baseline used $D = 128, 256, 512$ random features and bandwidth factor $g \in \{0.25, 1, 4\}$ with $\gamma = g/p$ for $p$ input features. The nonlinear neural correction used 512 features and $g = 1$ for three fixed maps. Tables S8–S14 and S19 give the corresponding gains, timing offsets, split seeds and replacement comparisons. We selected the 20 confirmation sessions before scoring them and used the same sessions for both penalty-selection procedures. The prespecified criteria for replication within this animal required positive geometry-baseline gains on at least 16 of 20 sessions and in both task means, a lower mean gain with position history, and a same-trial advantage on at least 16 sessions and in both task means.

### Calibration and shared geometry in simulation

The calibration generated 480 trials of 60 bins in eight conditions with 48 neural features. Velocity comprised a smooth condition-specific template and a trial deviation with an autoregressive coefficient of 0.85. Neural activity encoded the condition and time. The three regimes embedded the velocity deviation in neural activity at strengths 0, 0.8 and 1.8. All regimes used the same ten seeds. Median gain was approximately zero without the injected deviation and rose to 0.22451 and 0.37360 for the two nonzero strengths (Table S17).

In the shared-geometry simulation the baseline inputs were two continuous geometry variables and a five-term phase basis. The generating basis included phase, geometry, geometry by phase, the three quadratic geometry terms and their products with phase. Neural and velocity coefficients had scale $1.5/\sqrt{35}$. Neural noise was independent Gaussian noise with scale 0.20. Velocity noise had autoregressive coefficient 0.85 and innovation scale 0.20. Both baselines used the same data within each seed. All shared structure came from the measured baseline inputs. Table S21 reports each seed under the restricted and quadratic representations.

### Continuous reaching across animals and recording time

We selected ten dates per animal from the original Indy and Loco recordings. All identifiable M1 channels, including silent channels, entered the predictor. The target was the difference between successive 20 ms cursor-position averages divided by their timestamp difference. Neural and position inputs used bins $k-9,\ldots,k-5$. Task state used the current and previous target and elapsed time from the last change, evaluated at native sample $5(k-4)-1$. A target change required a displacement above the fixed $10^{-6}$ mm tolerance. We retained local bins 9–98 within each 2 s block. Whole blocks stayed in the same partition. We excluded blocks before the first observed target change, blocks with non-finite position or command state, and blocks with native sampling intervals outside 4 ms $\pm 2\,\mu$s. The split was defined on physical blocks before exclusions and required at least 100 eligible training and 40 eligible test blocks.

Task and position-history baselines had 5 and 15 linear predictors. Their primary quadratic forms had 20 and 135 columns. Matching allowed any other held-out block and ranked candidates by distance between training-standardized baseline trajectories. The models used seed 20261015. Tables S22–S24 give the selected recordings and the linear and quadratic estimates before capping elapsed time. The mean primary gain decreased from 0.2737 to 0.0572 in Indy and from 11.7338 to 0.0276 in Loco when position history entered the baseline. The Loco task mean included the 10 February 2017 session with $\Delta R^2 = 116.0740$. On that date 22.70% of test observations exceeded the training maximum for time since target change and contributed 99.69% of task-baseline squared error.

The post hoc cap was the largest elapsed time in the training subset for each nuisance fit. We applied it before quadratic expansion and used the outer-training cap to construct the replacement-matching inputs. The two model forms used identical observations. With this cap, mean early task and position-history gains were 0.2661 and 0.0454 in Indy and 0.1691 and 0.0289 in Loco (Tables S25 and S26). On 10 February the capped task baseline and full prediction had $R^2 = -1.2193$ and $-0.8073$, respectively. With uncapped quadratic terms and position history in Loco, neural gain was negative on one date ($-0.0100$). In Loco the capped task baseline had negative $R^2$ on four early dates and the corresponding full prediction had negative $R^2$ on three.

All ten Loco files contained the later interval. The selected Indy files lacked an interval of comparable duration. For each Loco date we kept the training observations and fitted coefficients fixed across the two test intervals. We verified agreement with the predictions for the first interval before scoring blocks 301–549, spanning $[602, 1100)$ s. Tables S27–S29 give the scores for all sixty later evaluations and the forty paired changes in error from capping elapsed time. The cap lowered full mean squared error on eight dates for task inputs and nine for position history. On 17 February 2017 it lowered the full task prediction's root mean squared error from 13,011.0 to 187.8 mm/s. The capped full prediction still had $R^2 = -102.5789$. Two capped gains were negative, $-0.02335$ for task inputs on 10 February and $-0.10100$ with position history on 14 February. Task-minus-history gain was positive on eight dates for linear models, eight for quadratic models and seven for capped quadratic models. Each of the three paired-date intervals for that contrast included zero. The changing range of elapsed time can account for part of the error. The later interval also permits changes in movement patterns and neural activity, whose contributions cannot be separated in these recordings.

### Analysis settings

**Table S1.** Ridge penalties, random seeds, resampling and replacement settings.

| Setting | Specification |
|---|---|
| Ridge regressions | Each of the three regressions selects among $0.1, 1, 10, 100, 10^3, 10^4, 10^5$ after training-subset standardization. |
| LINK partitions | Stratify by direction. Use 70/30 whole trials for outer training/testing, 80/20 for inner selection and five folds for the nuisance regressions. Benchmark recordings use their released training/validation labels. |
| Random seeds | Base seed 20260726. Add 21 for nuisance folds, 301 for neural regression selection, 401 for final nuisance selection, 501 for the trial bootstrap and 601 for permutations. Add $100 + k$ to the nuisance seed in fold $k$. |
| Nested selection | Within neural-regression selection fold $k$, add $1000 + 100k$ to the selection seed for nuisance cross-fitting, then add 50 for the fold's final nuisance selection. The inner holdout count is $\max(\lceil 0.2n \rceil, \text{number of conditions})$ when stratification is feasible. |
| LINK split sensitivity | Seeds 20260726, 20260727 and 20260728 vary the outer split and internal model selection together. |
| LINK confirmation | Ten center-out and ten random-target sessions. Both baselines use spiking-band power and seed 20260726. |
| Nonlinear and joint models | The joint ridge comparison selects separate penalties for baseline and neural predictors. The nonlinear baseline comparison uses 256 random Fourier features and kernel multipliers $0.25, 1, 4$. The neural correction uses 512 features, multiplier 1 and map seeds 20260802, 20260803 and 20260804. |
| Uncertainty | 500 whole-trial bootstrap samples per LINK session for primary, confirmation, timing and split analyses. For the model-family sensitivities, use 500 per LINK session and 1,000 per benchmark condition. LINK task contrasts use 10,000 paired-date resamples and all $2^9$ sign assignments. |
| Replacement | Among other held-out trials of the same condition, use the nearest complete baseline trajectory. Scale each feature on training trials and break ties by trial index. Subtract the source's own fitted baseline from its neural activity. |
| Simulation | 480 trials, 60 bins, eight conditions, 48 neural features and two velocity components. Template scale 1.0, autoregressive coefficient 0.85 and neural noise standard deviation 0.20. The behavioral deviation scales are 0.20, 0.25 and 0.35, with corresponding neural embedding strengths 0, 0.8 and 1.8. Each regime uses seeds 20260726–20260735. Within each seed, regimes share template, base deviation, weights and noise. The estimator uses the generator seed. |
| LINK timing | 20 ms bins, 160 ms of neural history ending before the velocity bin, 25 predicted bins and 100 ms of position history. The timing sensitivity uses 0, 40, 80 and 120 ms leads over the fixed 320–820 ms window. |
| Benchmark timing | 20 ms bins, 100 ms histories and an 80 ms lead for both neural and position inputs. |
| Scoring | Average squared error over test bins and both velocity components. The mean of their test variances normalizes $\Delta R^2$. Each session contributes equally to the group mean. |

**Table S2.** Data dimensions for the two LINK cohorts and the two benchmark recordings. CO denotes center-out and RD random-target.

| Dataset | Trials or blocks | Outer train/test | Neural channels/units | Time bins | Neural feature dimension | Primary context dimension |
|---|---|---|---|---|---|---|
| LINK, each session | 375 | 262/113 | 96 | 25 | 768 | 141 geometry; 151 with history |
| MC_Maze-medium | 250 | 188/62 | 152 | 40 | 760 | 258 template; 268 with history |
| MC_RTT | 1,080 | 810/270 | 130 | 21 | 650 | 2 target; 6 state; 14 with history |

### LINK baselines and paired task differences

**Table S3.** Spiking-band power in the LINK development sessions. Values are means across nine dates for each task. Five baseline specifications share the same trials and neural measurements.

| Context | Task | Context $R^2$ | Neural-only $R^2$ | Combined $R^2$ | Neural gain | Replacement gain |
|---|---|---|---|---|---|---|
| Phase | CO | 0.00838 | 0.3077 | 0.30958 | 0.30119 | -0.01087 |
| Phase | RD | 0.00546 | 0.3129 | 0.31794 | 0.31248 | 0.10498 |
| Direction × phase | CO | 0.40643 | 0.3077 | 0.47211 | 0.06567 | -0.04162 |
| Direction × phase | RD | 0.45239 | 0.3129 | 0.50403 | 0.05165 | -0.03488 |
| + geometry | CO | 0.46212 | 0.3077 | 0.50640 | 0.04428 | 0.01294 |
| + geometry | RD | 0.51755 | 0.3129 | 0.55079 | 0.03324 | -0.00818 |
| Geometry baseline | CO | 0.57022 | 0.3077 | 0.58956 | 0.01933 | -0.00237 |
| Geometry baseline | RD | 0.59994 | 0.3129 | 0.62079 | 0.02085 | -0.01191 |
| + position history | CO | 0.66167 | 0.3077 | 0.66933 | 0.00766 | -0.00154 |
| + position history | RD | 0.68388 | 0.3129 | 0.69388 | 0.01000 | -0.00614 |

**Table S4.** Difference in neural gain between LINK random-target and center-out sessions on the same nine dates. The sign-flip test enumerates all 512 assignments. Intervals resample paired dates.

| Context | Mean RD − CO gain | 95% date-bootstrap CI | Exact two-sided sign-flip $p$ |
|---|---|---|---|
| Phase | 0.01128 | [-0.03822, 0.05005] | 0.6719 |
| Direction × phase | -0.01403 | [-0.02681, -0.00315] | 0.0430 |
| + geometry | -0.01104 | [-0.02163, -0.00200] | 0.0508 |
| Geometry baseline | 0.00152 | [-0.00356, 0.00764] | 0.6992 |
| + position history | 0.00234 | [-0.00119, 0.00672] | 0.3672 |

### Benchmark recordings

**Table S5.** Exploratory benchmark baselines with penalties selected on a single validation split. Substituted neural activity was residualized against the target trial's inputs in this comparison.

| Dataset | Context | Context $R^2$ | Neural-only $R^2$ | Neural gain | 95% trial-bootstrap CI | Replacement gain |
|---|---|---|---|---|---|---|
| MC_Maze | template | 0.9053 | 0.4812 | 0.00011 | [-0.00143, 0.00148] | -0.00277 |
| MC_Maze | + lag-matched history | 0.9248 | 0.4812 | 0.00019 | [-0.00099, 0.00135] | -0.00273 |
| MC_RTT | target only | 0.1024 | 0.3683 | 0.27627 | [0.24195, 0.31177] | -0.17708 |
| MC_RTT | task state | 0.2088 | 0.3683 | 0.24090 | [0.21152, 0.27296] | 0.01554 |
| MC_RTT | + lag-matched history | 0.6031 | 0.3683 | 0.06846 | [0.05448, 0.08434] | -0.01554 |

**Table S6.** Neural gain in two benchmark recordings with the fully nested estimator. Intervals resample held-out trials or blocks within condition. The neural random-feature model uses map seed 20260802.

| Dataset | Model | Context $R^2$ | Neural gain (95% CI) | Replacement | MSE reduction |
|---|---|---|---|---|---|
| MC_Maze | linear residual | 0.90533 | 0.000513 (0.000257, 0.000757) | -0.000037 | 0.54% |
| MC_Maze | neural RFF | 0.90533 | 0.000045 (—) | 0.000043 | 0.05% |
| MC_RTT | linear residual | 0.60308 | 0.068460 (0.053709, 0.084777) | -0.022973 | 17.25% |
| MC_RTT | neural RFF | 0.60308 | 0.016851 (—) | -0.005527 | 4.25% |

### Neural signals, baseline forms and timing

**Table S7.** LINK neural gain with spiking-band power (SBP) and threshold-crossing rate (TCR) recorded on the same electrodes. The paired-date sign-flip tests are descriptive.

| Signal | Context | CO gain | RD gain | RD − CO exact sign-flip $p$ |
|---|---|---|---|---|
| SBP | geometry baseline | 0.01933 | 0.02085 | 0.6992 |
| SBP | + position history | 0.00766 | 0.01000 | 0.3672 |
| TCR | geometry baseline | 0.00992 | 0.01263 | 0.2773 |
| TCR | + position history | 0.00329 | 0.00539 | 0.1250 |

**Table S8.** Quadratic geometry in LINK. The additional terms expand the geometry baseline using the same measured inputs. Penalties were selected on a single validation split.

| Signal | Context | CO gain | RD gain | Mean RD − CO | 95% date-bootstrap CI | Exact $p$ |
|---|---|---|---|---|---|---|
| SBP | quadratic geometry | 0.01599 | 0.01958 | 0.00358 | [-0.00092, 0.00900] | 0.2266 |
| SBP | quadratic geometry + history | 0.00540 | 0.00950 | 0.00410 | [0.00115, 0.00820] | 0.0234 |
| TCR | quadratic geometry | 0.00717 | 0.01235 | 0.00519 | [0.00122, 0.00925] | 0.0625 |

| Signal | Context | CO gain | RD gain | Mean RD − CO | 95% date-bootstrap CI | Exact $p$ |
|---|---|---|---|---|---|---|
| TCR | quadratic geometry + history | 0.00164 | 0.00538 | 0.00374 | [0.00142, 0.00653] | 0.0117 |

**Table S9.** Relative timing of neural and velocity measurements across a fixed behavioral window in LINK. Each offset uses the same penalty-selection procedure as Table S8.

| Neural/output lead | CO gain | RD gain | Mean RD − CO |
|---|---|---|---|
| 0 ms | 0.00410 | 0.00615 | 0.00205 |
| 40 ms | 0.00143 | 0.00008 | -0.00135 |
| 80 ms | 0.00071 | -0.00082 | -0.00153 |
| 120 ms | 0.00068 | 0.00067 | -0.00001 |

**Table S10.** Variation across three train/test splits in LINK. Within each training set, one validation split selected the penalties. The source data also give the estimates with fully nested penalty selection.

| Split seed | CO geometry gain | RD geometry gain | RD − CO 95% CI |
|---|---|---|---|
| 20260726 | 0.01974 | 0.02045 | [-0.00402, 0.00684] |
| 20260727 | 0.01833 | 0.02193 | [-0.00141, 0.00896] |
| 20260728 | 0.01622 | 0.01988 | [-0.00140, 0.00905] |

**Table S11.** Direct concatenation of LINK baseline and neural inputs with a shared ridge penalty. A single validation split selected the common penalty.

| Signal/context | CO direct gain | RD direct gain |
|---|---|---|
| SBP geometry baseline | -0.0180 | -0.0075 |
| SBP + position history | -0.0260 | -0.0203 |
| TCR geometry baseline | -0.0402 | -0.0236 |
| TCR + position history | -0.0383 | -0.0303 |

**Table S12.** Sensitivity to model form. We compared residual regression, separate penalties for baseline and neural inputs, and a nonlinear baseline. LINK entries average 18 sessions. Each benchmark dataset contributes one recording.

| Dataset/signal | Context | Residual-neural | Separate block | Nonlinear context | Source replacement |
|---|---|---|---|---|---|
| LINK SBP | geometry baseline | 0.02010 | 0.01924 | 0.01733 | -0.00735 |
| LINK SBP | + position history | 0.00891 | 0.00855 | 0.00663 | -0.00387 |
| LINK TCR | geometry baseline | 0.01158 | 0.01102 | 0.00949 | -0.00865 |
| LINK TCR | + position history | 0.00424 | 0.00410 | 0.00274 | -0.00458 |
| MC_Maze | task template | 0.00011 | 0.00046 | -0.00024 | -0.00280 |
| MC_RTT | + lag-matched history | 0.06846 | 0.06902 | 0.04619 | -0.02297 |

**Table S13.** Sensitivity to the number of random Fourier features and the random seed for the baseline features. Values average the sessions in each named group.

| Endpoint | Baseline | Seed 2 | Seed 3 | $D = 128$ | $D = 512$ |
|---|---|---|---|---|---|
| LINK SBP, CO | 0.01461 | 0.01545 | 0.01480 | 0.01651 | 0.01470 |
| LINK SBP, RD | 0.02006 | 0.01915 | 0.02030 | 0.01995 | 0.01951 |
| LINK TCR, CO | 0.00686 | 0.00734 | 0.00715 | 0.00766 | 0.00637 |

| Endpoint | Baseline | Seed 2 | Seed 3 | $D = 128$ | $D = 512$ |
|---|---|---|---|---|---|
| LINK TCR, RD | 0.01212 | 0.01251 | 0.01209 | 0.01211 | 0.01183 |
| MC_Maze | -0.00024 | 0.00045 | 0.00050 | -0.00021 | 0.00059 |
| MC_RTT | 0.04619 | 0.04571 | 0.04764 | 0.05708 | 0.04077 |

**Table S14.** Gain for each of the five nearest source trials in LINK. Rank one has the smallest distance between baseline trajectories.

| Signal/task | Exact | Rank 1 | Rank 2 | Rank 3 | Rank 4 | Rank 5 | Pool mean |
|---|---|---|---|---|---|---|---|
| SBP, CO | 0.01974 | -0.00250 | -0.00659 | -0.00692 | -0.00819 | -0.00996 | -0.00683 |
| SBP, RD | 0.02045 | -0.01220 | -0.01091 | -0.01413 | -0.01526 | -0.01404 | -0.01330 |
| TCR, CO | 0.01073 | -0.00575 | -0.00833 | -0.00784 | -0.00981 | -0.01118 | -0.00858 |
| TCR, RD | 0.01243 | -0.01127 | -0.01189 | -0.01210 | -0.01139 | -0.01238 | -0.01186 |

### Cohorts and simulation controls

**Table S15.** Analysis sets and their units of comparison.

| Analysis set | Sampling unit | Comparison |
|---|---|---|
| LINK development | Nine recording dates | Nine pairs of center-out and random-target sessions in one animal. |
| LINK confirmation | Twenty sessions | Ten sessions per task selected by date and asset rank before their first analysis. |
| Benchmark recordings | One session per task | MC_Maze and MC_RTT use release-defined partitions. |
| Reaching, first interval | Ten dates per animal | Indy and Loco models use separate fits for each recording. |
| Reaching, later interval | Same ten Loco dates | Fixed models applied to 249 subsequent blocks per date. |
| Simulation | Ten seeds per comparison | Both baselines use the same generated observations within each seed. |

**Table S16.** LINK development estimates with the fully nested estimator. Each interval resamples the nine dates from the same animal. Percentage reduction uses the baseline mean squared error as its denominator.

| Signal | Context | Mean gain (95% date CI) | Replacement | MSE reduction |
|---|---|---|---|---|
| SBP | geometry baseline | 0.02009 (0.01586, 0.02404) | -0.00714 | 4.92% |
| SBP | + position history | 0.00883 (0.00628, 0.01097) | -0.00384 | 2.75% |
| TCR | geometry baseline | 0.01127 (0.00665, 0.01584) | -0.00830 | 2.75% |
| TCR | + position history | 0.00434 (0.00245, 0.00633) | -0.00488 | 1.37% |

**Table S17.** Calibration with ten matched seeds per regime. Parentheses give the observed range across seeds.

| Regime | Median gain (seed range) | Mean replacement | Mean permutation |
|---|---|---|---|
| Template only | -0.000002 (-0.000059, 0.000025) | -0.000019 | -0.000013 |
| Weak trial signal | 0.22451 (0.21215, 0.24304) | -0.20897 | -0.21933 |
| Strong trial signal | 0.37360 (0.35383, 0.40353) | -0.34859 | -0.36631 |

**Table S18.** Twenty LINK confirmation sessions. Each pair of columns gives the fully nested same-trial gain and the gain after trial replacement for one baseline.

| Date | Task | Geometry | Geometry repl. | + history | History repl. |
|---|---|---|---|---|---|
| 2020-09-04 | RD | 0.04001 | -0.02781 | 0.01130 | -0.02008 |
| 2020-09-24 | CO | 0.02186 | -0.00018 | 0.01058 | -0.00036 |
| 2020-12-11 | CO | 0.01049 | 0.00608 | 0.00774 | 0.00466 |
| 2021-03-23 | CO | 0.01497 | -0.00692 | 0.00436 | -0.00745 |
| 2021-04-28 | RD | 0.02400 | -0.01016 | 0.01291 | -0.00793 |
| 2021-06-01 | CO | 0.01503 | -0.00830 | 0.00396 | -0.00422 |
| 2021-07-01 | RD | 0.02151 | -0.00856 | 0.01045 | -0.00569 |
| 2021-08-04 | RD | 0.00809 | -0.00878 | 0.00390 | -0.00495 |
| 2021-08-05 | CO | 0.01457 | -0.00385 | 0.00290 | 0.00136 |
| 2021-09-15 | CO | 0.02289 | 0.00166 | 0.01017 | 0.00303 |
| 2021-11-16 | CO | 0.02879 | -0.00434 | 0.01275 | -0.00096 |
| 2022-02-10 | RD | 0.01349 | -0.01204 | 0.00486 | -0.00900 |
| 2022-02-28 | CO | 0.03433 | 0.00517 | 0.00510 | -0.00007 |
| 2022-04-27 | CO | 0.01654 | -0.00424 | 0.00530 | -0.00081 |
| 2022-05-11 | RD | 0.02752 | -0.00494 | 0.01206 | -0.00701 |
| 2022-08-25 | RD | 0.03650 | -0.01573 | 0.01271 | -0.00993 |
| 2022-09-02 | CO | 0.01734 | -0.01169 | 0.00410 | -0.00706 |
| 2023-01-24 | RD | 0.02576 | -0.02042 | 0.00935 | -0.01040 |
| 2023-03-10 | RD | 0.00593 | -0.00638 | -0.00128 | -0.00499 |
| 2023-04-26 | RD | 0.01845 | -0.01022 | 0.00770 | -0.00781 |

**Table S19.** Neural random Fourier feature estimates across three fixed maps. LINK geometry values average nine dates per task. The position-history values average all 18 sessions. Each benchmark contributes one recording.

| Map seed | LINK CO geom. | LINK RD geom. | LINK history | MC_Maze | MC_RTT |
|---|---|---|---|---|---|
| 20260802 | 0.004315 | 0.003205 | 0.000947 | 0.000045 | 0.016851 |
| 20260803 | 0.003914 | 0.003904 | 0.001254 | 0.000077 | 0.015012 |
| 20260804 | 0.003514 | 0.002863 | 0.000988 | 0.000139 | 0.014908 |

**Table S20.** Inputs and time references for the velocity predictions.

| Dataset | Inputs preceding velocity | Time reference |
|---|---|---|
| LINK | 160 ms of SBP or TCR. The position-history baseline adds 100 ms of finger position. | Histories end at bin $k-1$. Velocity uses positions in bins $k$ and $k+1$. |
| MC_Maze | 100 ms of spikes with an 80 ms lead. | The window starts at measured movement onset. |
| MC_RTT | Spikes and measured cursor history have a common 80 ms lead. | The block's mean error direction defines retrospective matching strata. |
| Indy and Loco reaching | 100 ms of M1 spikes and, for the position-history baseline, 100 ms of cursor position. | Both histories end 80 ms before the target bin. Task state is sampled before that boundary. |

**Table S21.** Shared-geometry simulation. The linear and quadratic baselines use the same observations and split for each seed. Repl. denotes gain after trial replacement.

| Seed | Linear gain | Linear repl. | Quadratic gain | Quadratic repl. |
|---|---|---|---|---|
| 20260726 | 0.58506 | 0.34764 | 0.00000417 | 0.00000556 |
| 20260727 | 0.46123 | 0.38927 | -0.00004928 | -0.00000906 |
| 20260728 | 0.71554 | 0.50915 | -0.00001281 | -0.00001556 |
| 20260729 | 0.49619 | 0.34148 | -0.00000665 | -0.00000551 |

| Seed | Linear gain | Linear repl. | Quadratic gain | Quadratic repl. |
|---|---|---|---|---|
| 20260730 | 0.55388 | 0.40221 | -0.00000547 | -0.00000006 |
| 20260731 | 0.45775 | 0.34050 | 0.00000092 | 0.00000340 |
| 20260732 | 0.63628 | 0.44059 | 0.00000096 | 0.00000502 |
| 20260733 | 0.58459 | 0.29309 | -0.00000959 | -0.00000486 |
| 20260734 | 0.71354 | 0.50031 | 0.00001186 | 0.00000646 |
| 20260735 | 0.65305 | 0.51412 | -0.00000144 | 0.00000050 |

### Early continuous-reaching evaluation

**Table S22.** Selected reaching recordings. Duration is the first analysis interval. Train and test counts are eligible two-second blocks.

| Animal | File date/suffix | Duration (s) | M1 | Train | Test |
|---|---|---|---|---|---|
| Indy | 20160407/02 | 600.000 | 96 | 207 | 89 |
| Indy | 20160419/01 | 524.004 | 96 | 182 | 78 |
| Indy | 20160624/03 | 500.000 | 96 | 172 | 74 |
| Indy | 20160916/01 | 451.828 | 96 | 156 | 67 |
| Indy | 20160930/02 | 460.788 | 96 | 160 | 68 |
| Indy | 20161011/03 | 600.000 | 96 | 209 | 89 |
| Indy | 20161024/03 | 472.352 | 96 | 164 | 70 |
| Indy | 20161027/03 | 578.960 | 96 | 201 | 86 |
| Indy | 20161220/02 | 576.328 | 96 | 200 | 86 |
| Indy | 20170131/02 | 600.000 | 96 | 209 | 89 |
| Loco | 20170210/03 | 600.000 | 96 | 209 | 89 |
| Loco | 20170213/02 | 600.000 | 96 | 209 | 89 |
| Loco | 20170214/02 | 600.000 | 96 | 208 | 89 |
| Loco | 20170215/02 | 600.000 | 96 | 209 | 89 |
| Loco | 20170216/02 | 600.000 | 96 | 208 | 89 |
| Loco | 20170217/02 | 600.000 | 96 | 206 | 89 |
| Loco | 20170227/04 | 600.000 | 96 | 209 | 89 |
| Loco | 20170228/02 | 600.000 | 96 | 209 | 89 |
| Loco | 20170301/05 | 600.000 | 96 | 209 | 89 |
| Loco | 20170302/02 | 600.000 | 96 | 208 | 89 |

**Table S23.** Early Indy interval with uncapped linear and quadratic models. T and H denote task and position-history inputs. L and Q denote linear and quadratic terms. C/F gives baseline and full root mean squared error in mm/s. Gain intervals resample held-out blocks.

| Date | Basis | C $R^2$ | Gain [95% interval] | Replace | RMSE C/F | MSE ↓ (%) |
|---|---|---|---|---|---|---|
| 20160407 | TL | 0.160 | 0.3129 [0.2830, 0.3423] | -0.1673 | 52.2/41.3 | 37.26 |
| 20160407 | TQ | 0.206 | 0.2703 [0.2336, 0.3085] | -0.1556 | 50.7/41.2 | 34.06 |
| 20160407 | HL | 0.681 | 0.0390 [0.0295, 0.0501] | -0.0255 | 32.2/30.1 | 12.22 |
| 20160407 | HQ | 0.784 | 0.0148 [0.0092, 0.0223] | -0.0161 | 26.5/25.6 | 6.84 |
| 20160419 | TL | 0.111 | 0.2775 [0.2358, 0.3297] | -0.0674 | 58.4/48.4 | 31.23 |
| 20160419 | TQ | -0.869 | 0.7045 [0.3716, 1.1929] | 0.1842 | 84.6/66.8 | 37.69 |
| 20160419 | HL | 0.646 | 0.0498 [0.0352, 0.0701] | -0.0114 | 36.8/34.1 | 14.09 |
| 20160419 | HQ | 0.544 | 0.0259 [-0.0106, 0.0630] | -0.0121 | 41.8/40.6 | 5.69 |
| 20160624 | TL | 0.216 | 0.2758 [0.2317, 0.3291] | -0.1307 | 55.4/44.6 | 35.18 |
| 20160624 | TQ | 0.261 | 0.2412 [0.2047, 0.2815] | -0.1357 | 53.8/44.1 | 32.62 |
| 20160624 | HL | 0.567 | 0.0864 [0.0663, 0.1156] | 0.0040 | 41.2/36.8 | 19.95 |
| 20160624 | HQ | -0.308 | 0.1683 [0.1004, 0.2683] | 0.0864 | 71.5/66.8 | 12.87 |
| 20160916 | TL | 0.226 | 0.2782 [0.2523, 0.3046] | -0.1668 | 68.7/55.0 | 35.93 |
| 20160916 | TQ | 0.294 | 0.2369 [0.2110, 0.2631] | -0.1480 | 65.6/53.5 | 33.55 |
| 20160916 | HL | 0.646 | 0.0653 [0.0520, 0.0798] | -0.0577 | 46.5/42.0 | 18.43 |
| 20160916 | HQ | 0.736 | 0.0453 [0.0333, 0.0622] | -0.0240 | 40.1/36.5 | 17.19 |

| Date | Basis | C $R^2$ | Gain [95% interval] | Replace | RMSE C/F | MSE ↓ (%) |
|---|---|---|---|---|---|---|
| 20160930 | TL | 0.218 | 0.2175 [0.1771, 0.2452] | -0.1103 | 63.5/53.9 | 27.82 |
| 20160930 | TQ | 0.208 | 0.2378 [0.2089, 0.2628] | -0.1177 | 63.9/53.4 | 30.02 |
| 20160930 | HL | 0.657 | 0.0410 [0.0316, 0.0492] | -0.0213 | 42.1/39.5 | 11.95 |
| 20160930 | HQ | 0.417 | 0.1180 [0.1053, 0.1322] | -0.0303 | 54.8/48.9 | 20.25 |
| 20161011 | TL | 0.221 | 0.2831 [0.2633, 0.3047] | -0.1223 | 67.3/53.7 | 36.36 |
| 20161011 | TQ | 0.313 | 0.2265 [0.2077, 0.2457] | -0.1519 | 63.2/51.7 | 32.96 |
| 20161011 | HL | 0.662 | 0.0667 [0.0586, 0.0764] | -0.0295 | 44.3/39.7 | 19.72 |
| 20161011 | HQ | 0.717 | 0.0267 [0.0239, 0.0299] | -0.0028 | 40.5/38.6 | 9.45 |
| 20161024 | TL | 0.208 | 0.2835 [0.2531, 0.3135] | -0.0435 | 73.6/59.0 | 35.79 |
| 20161024 | TQ | 0.335 | 0.2072 [0.1788, 0.2348] | -0.1019 | 67.4/55.9 | 31.17 |
| 20161024 | HL | 0.657 | 0.0719 [0.0603, 0.0850] | -0.0491 | 48.4/43.1 | 20.96 |
| 20161024 | HQ | 0.775 | 0.0263 [0.0209, 0.0326] | -0.0112 | 39.2/36.8 | 11.72 |
| 20161027 | TL | 0.206 | 0.2979 [0.2623, 0.3347] | -0.2667 | 60.5/47.8 | 37.53 |
| 20161027 | TQ | 0.292 | 0.2551 [0.2222, 0.2896] | -0.2648 | 57.2/45.7 | 36.02 |
| 20161027 | HL | 0.632 | 0.0850 [0.0707, 0.0993] | -0.0649 | 41.2/36.1 | 23.11 |
| 20161027 | HQ | 0.549 | 0.0974 [0.0812, 0.1165] | -0.0729 | 45.6/40.4 | 21.62 |
| 20161220 | TL | 0.227 | 0.2301 [0.2054, 0.2538] | -0.1075 | 65.4/54.8 | 29.75 |
| 20161220 | TQ | 0.338 | 0.1671 [0.1472, 0.1854] | -0.1390 | 60.5/52.3 | 25.26 |
| 20161220 | HL | 0.658 | 0.0450 [0.0365, 0.0538] | -0.0541 | 43.5/40.5 | 13.14 |
| 20161220 | HQ | 0.771 | 0.0193 [0.0144, 0.0250] | -0.0128 | 35.5/34.0 | 8.43 |
| 20170131 | TL | 0.234 | 0.2383 [0.2151, 0.2678] | -0.1332 | 59.3/49.2 | 31.13 |
| 20170131 | TQ | 0.339 | 0.1903 [0.1707, 0.2147] | -0.1476 | 55.1/46.5 | 28.81 |
| 20170131 | HL | 0.668 | 0.0616 [0.0545, 0.0706] | -0.0251 | 39.0/35.2 | 18.56 |
| 20170131 | HQ | 0.745 | 0.0297 [0.0244, 0.0358] | -0.0173 | 34.2/32.2 | 11.62 |

**Table S24.** Early Loco interval with uncapped linear and quadratic models. T/H denotes task/position history and L/Q denotes linear/quadratic. C/F gives baseline/full root mean squared error in mm/s. Gain intervals resample held-out blocks.

| Date | Basis | C $R^2$ | Gain [95% interval] | Replace | RMSE C/F | MSE ↓ (%) |
|---|---|---|---|---|---|---|
| 20170210 | TL | -0.830 | 0.6832 [0.4370, 1.0466] | 0.3725 | 45.9/36.3 | 37.35 |
| 20170210 | TQ | -330.076 | 116.0740 [47.7750, 213.9206] | 99.4938 | 617.0/497.2 | 35.06 |
| 20170210 | HL | 0.588 | 0.0279 [0.0153, 0.0424] | -0.0173 | 21.8/21.0 | 6.77 |
| 20170210 | HQ | -1.888 | 0.0262 [-0.0949, 0.1438] | -0.0228 | 57.6/57.4 | 0.91 |
| 20170213 | TL | 0.075 | 0.1699 [0.1318, 0.2024] | -0.2244 | 34.8/31.4 | 18.36 |
| 20170213 | TQ | 0.104 | 0.1475 [0.1057, 0.1811] | -0.2052 | 34.2/31.3 | 16.46 |
| 20170213 | HL | 0.469 | 0.0394 [0.0250, 0.0523] | -0.0343 | 26.4/25.4 | 7.41 |
| 20170213 | HQ | 0.475 | 0.0353 [0.0240, 0.0479] | -0.0286 | 26.2/25.3 | 6.72 |
| 20170214 | TL | 0.101 | 0.1378 [0.1115, 0.1604] | -0.1048 | 28.4/26.1 | 15.34 |
| 20170214 | TQ | 0.075 | 0.1142 [0.0852, 0.1395] | -0.1111 | 28.8/27.0 | 12.36 |
| 20170214 | HL | 0.507 | 0.0201 [0.0088, 0.0328] | -0.0206 | 21.0/20.6 | 4.08 |
| 20170214 | HQ | 0.531 | -0.0100 [-0.0408, 0.0091] | -0.0477 | 20.5/20.7 | -2.13 |
| 20170215 | TL | 0.115 | 0.0801 [0.0564, 0.1021] | -0.0998 | 30.6/29.2 | 9.04 |
| 20170215 | TQ | -0.050 | 0.1290 [0.0943, 0.1793] | -0.0349 | 33.3/31.2 | 12.29 |
| 20170215 | HL | 0.514 | 0.0062 [-0.0013, 0.0136] | -0.0250 | 22.7/22.5 | 1.28 |
| 20170215 | HQ | -0.072 | 0.1302 [0.0460, 0.2854] | 0.0935 | 33.7/31.6 | 12.15 |
| 20170216 | TL | 0.119 | 0.0950 [0.0668, 0.1204] | -0.0935 | 34.0/32.1 | 10.79 |
| 20170216 | TQ | 0.142 | 0.0848 [0.0576, 0.1106] | -0.0878 | 33.5/31.8 | 9.89 |
| 20170216 | HL | 0.588 | 0.0096 [0.0033, 0.0165] | -0.0138 | 23.2/23.0 | 2.33 |
| 20170216 | HQ | 0.510 | 0.0249 [0.0065, 0.0492] | 0.0041 | 25.3/24.7 | 5.08 |
| 20170217 | TL | 0.108 | 0.1315 [0.1055, 0.1554] | -0.0907 | 35.3/32.6 | 14.74 |
| 20170217 | TQ | 0.125 | 0.1188 [0.0954, 0.1429] | -0.0781 | 34.9/32.5 | 13.58 |
| 20170217 | HL | 0.508 | 0.0236 [0.0151, 0.0328] | -0.0259 | 26.2/25.5 | 4.80 |
| 20170217 | HQ | 0.556 | 0.0159 [0.0121, 0.0205] | -0.0006 | 24.9/24.4 | 3.59 |
| 20170227 | TL | 0.189 | 0.0987 [0.0365, 0.1441] | -0.2327 | 24.4/22.9 | 12.16 |
| 20170227 | TQ | 0.183 | 0.0891 [0.0284, 0.1337] | -0.2218 | 24.5/23.1 | 10.90 |
| 20170227 | HL | 0.656 | 0.0107 [0.0044, 0.0172] | -0.0224 | 15.9/15.6 | 3.11 |
| 20170227 | HQ | 0.648 | 0.0052 [-0.0039, 0.0137] | -0.0258 | 16.1/15.9 | 1.49 |

| Date | Basis | C $R^2$ | Gain [95% interval] | Replace | RMSE C/F | MSE ↓ (%) |
|---|---|---|---|---|---|---|
| 20170228 | TL | 0.137 | 0.1718 [0.1372, 0.2054] | -0.1431 | 33.9/30.4 | 19.92 |
| 20170228 | TQ | -0.745 | 0.1117 [0.0393, 0.1697] | -0.2228 | 48.3/46.7 | 6.40 |
| 20170228 | HL | 0.662 | 0.0173 [0.0122, 0.0231] | -0.0120 | 21.3/20.7 | 5.12 |
| 20170228 | HQ | 0.645 | 0.0154 [0.0097, 0.0220] | -0.0059 | 21.8/21.3 | 4.34 |
| 20170301 | TL | 0.128 | 0.1504 [0.1139, 0.1903] | -0.1443 | 35.0/31.8 | 17.24 |
| 20170301 | TQ | -0.674 | 0.3613 [0.1762, 0.6618] | -0.0590 | 48.5/42.9 | 21.58 |
| 20170301 | HL | 0.626 | 0.0233 [0.0140, 0.0316] | -0.0174 | 22.9/22.2 | 6.24 |
| 20170301 | HQ | 0.482 | 0.0229 [0.0069, 0.0427] | -0.0202 | 27.0/26.4 | 4.42 |
| 20170302 | TL | 0.159 | 0.1114 [0.0595, 0.1541] | -0.0613 | 25.0/23.3 | 13.24 |
| 20170302 | TQ | 0.159 | 0.1078 [0.0672, 0.1456] | -0.0667 | 25.0/23.3 | 12.82 |
| 20170302 | HL | 0.655 | 0.0022 [-0.0121, 0.0117] | -0.0150 | 16.0/15.9 | 0.63 |
| 20170302 | HQ | 0.686 | 0.0096 [0.0003, 0.0174] | -0.0119 | 15.3/15.0 | 3.06 |

### Elapsed time capped within training folds

**Table S25.** Early Indy interval with elapsed time capped at its training maximum before quadratic expansion. BT and BH denote capped task and position-history models. C/F gives baseline/full root mean squared error in mm/s.

| Date | Basis | C $R^2$ | Gain [95% interval] | Replace | RMSE C/F | MSE ↓ (%) |
|---|---|---|---|---|---|---|
| 20160407 | BT | 0.219 | 0.2641 [0.2310, 0.2968] | -0.1576 | 50.3/41.0 | 33.80 |
| 20160407 | BH | 0.790 | 0.0144 [0.0090, 0.0211] | -0.0158 | 26.1/25.2 | 6.86 |
| 20160419 | BT | -0.869 | 0.7044 [0.3716, 1.1929] | 0.1842 | 84.6/66.8 | 37.69 |
| 20160419 | BH | 0.544 | 0.0259 [-0.0106, 0.0630] | -0.0121 | 41.8/40.6 | 5.69 |
| 20160624 | BT | 0.261 | 0.2412 [0.2047, 0.2815] | -0.1357 | 53.8/44.1 | 32.62 |
| 20160624 | BH | -0.307 | 0.1683 [0.1004, 0.2682] | 0.0864 | 71.5/66.7 | 12.87 |
| 20160916 | BT | 0.311 | 0.2274 [0.2010, 0.2552] | -0.1602 | 64.8/53.1 | 32.99 |
| 20160916 | BH | 0.736 | 0.0455 [0.0334, 0.0625] | -0.0241 | 40.1/36.5 | 17.25 |
| 20160930 | BT | 0.271 | 0.1767 [0.1411, 0.2029] | -0.1764 | 61.3/53.3 | 24.25 |
| 20160930 | BH | 0.716 | 0.0004 [-0.0146, 0.0117] | -0.0697 | 38.2/38.2 | 0.13 |
| 20161011 | BT | 0.313 | 0.2265 [0.2077, 0.2457] | -0.1519 | 63.2/51.7 | 32.96 |
| 20161011 | BH | 0.717 | 0.0267 [0.0239, 0.0299] | -0.0028 | 40.5/38.6 | 9.45 |
| 20161024 | BT | 0.335 | 0.2072 [0.1788, 0.2348] | -0.1019 | 67.4/55.9 | 31.17 |
| 20161024 | BH | 0.775 | 0.0263 [0.0209, 0.0326] | -0.0112 | 39.2/36.8 | 11.72 |
| 20161027 | BT | 0.292 | 0.2553 [0.2226, 0.2897] | -0.2627 | 57.2/45.7 | 36.05 |
| 20161027 | BH | 0.549 | 0.0974 [0.0811, 0.1165] | -0.0728 | 45.6/40.4 | 21.61 |
| 20161220 | BT | 0.338 | 0.1673 [0.1475, 0.1854] | -0.1388 | 60.5/52.3 | 25.28 |
| 20161220 | BH | 0.771 | 0.0193 [0.0145, 0.0251] | -0.0128 | 35.5/34.0 | 8.45 |
| 20170131 | BT | 0.339 | 0.1906 [0.1711, 0.2149] | -0.1469 | 55.1/46.5 | 28.85 |
| 20170131 | BH | 0.745 | 0.0297 [0.0244, 0.0357] | -0.0172 | 34.2/32.2 | 11.62 |

**Table S26.** Early Loco interval with elapsed time capped at its training maximum before quadratic expansion. BT and BH denote capped task and position-history models. C/F gives baseline/full root mean squared error in mm/s.

| Date | Basis | C $R^2$ | Gain [95% interval] | Replace | RMSE C/F | MSE ↓ (%) |
|---|---|---|---|---|---|---|
| 20170210 | BT | -1.219 | 0.4120 [0.2958, 0.5772] | 0.0687 | 50.5/45.6 | 18.57 |
| 20170210 | BH | 0.498 | 0.0026 [-0.0109, 0.0161] | -0.0366 | 24.0/24.0 | 0.52 |
| 20170213 | BT | 0.109 | 0.1485 [0.1068, 0.1817] | -0.2129 | 34.1/31.2 | 16.67 |
| 20170213 | BH | 0.478 | 0.0354 [0.0243, 0.0481] | -0.0282 | 26.1/25.2 | 6.78 |
| 20170214 | BT | 0.151 | 0.1125 [0.0853, 0.1364] | -0.1099 | 27.6/25.7 | 13.24 |
| 20170214 | BH | 0.563 | 0.0119 [0.0010, 0.0235] | -0.0300 | 19.8/19.5 | 2.72 |
| 20170215 | BT | -0.050 | 0.1290 [0.0943, 0.1793] | -0.0349 | 33.3/31.2 | 12.29 |
| 20170215 | BH | -0.072 | 0.1302 [0.0460, 0.2853] | 0.0935 | 33.7/31.6 | 12.15 |
| 20170216 | BT | 0.142 | 0.0848 [0.0576, 0.1105] | -0.0877 | 33.5/31.8 | 9.88 |

| Date | Basis | C $R^2$ | Gain [95% interval] | Replace | RMSE C/F | MSE ↓ (%) |
|---|---|---|---|---|---|---|
| 20170216 | BH | 0.510 | 0.0241 [0.0064, 0.0478] | 0.0037 | 25.3/24.7 | 4.93 |
| 20170217 | BT | 0.125 | 0.1189 [0.0956, 0.1430] | -0.0781 | 34.9/32.5 | 13.59 |
| 20170217 | BH | 0.556 | 0.0199 [0.0118, 0.0286] | -0.0214 | 24.9/24.3 | 4.47 |
| 20170227 | BT | 0.182 | 0.0894 [0.0284, 0.1338] | -0.2237 | 24.5/23.1 | 10.94 |
| 20170227 | BH | 0.650 | 0.0058 [-0.0029, 0.0141] | -0.0244 | 16.0/15.9 | 1.66 |
| 20170228 | BT | -0.745 | 0.1118 [0.0393, 0.1697] | -0.2229 | 48.3/46.7 | 6.40 |
| 20170228 | BH | 0.645 | 0.0155 [0.0098, 0.0221] | -0.0058 | 21.8/21.3 | 4.37 |
| 20170301 | BT | -0.674 | 0.3759 [0.1795, 0.6942] | -0.0503 | 48.5/42.7 | 22.45 |
| 20170301 | BH | 0.482 | 0.0340 [0.0191, 0.0542] | -0.0025 | 27.0/26.1 | 6.57 |
| 20170302 | BT | 0.159 | 0.1080 [0.0673, 0.1459] | -0.0657 | 25.0/23.3 | 12.84 |
| 20170302 | BH | 0.686 | 0.0096 [0.0003, 0.0174] | -0.0119 | 15.3/15.0 | 3.06 |

### Later Loco evaluation

**Table S27.** Later Loco interval with task inputs. L, Q and B denote linear, quadratic and capped quadratic models. C/F gives baseline/full root mean squared error in mm/s. Gain intervals resample held-out blocks.

| Date | Basis | C $R^2$ | Gain [95% interval] | Replace | RMSE C/F | F $R^2$ |
|---|---|---|---|---|---|---|
| 20170210 | L | -19.014 | 10.6041 [7.7620, 14.5426] | 10.3338 | 124.0/85.0 | -8.410 |
| 20170210 | Q | -39556.9 | 13787.8 [9655.3, 19220.1] | 13654.2 | 5511.1/4448.1 | -25769.1 |
| 20170210 | B | -4.057 | -0.0233 [-0.0919, 0.0372] | -0.4810 | 62.3/62.5 | -4.080 |
| 20170213 | L | 0.044 | 0.1232 [0.0899, 0.1528] | -0.2806 | 30.9/28.9 | 0.167 |
| 20170213 | Q | -0.186 | 0.0723 [0.0280, 0.1109] | -0.2881 | 34.4/33.4 | -0.114 |
| 20170213 | B | -0.078 | 0.0778 [0.0384, 0.1139] | -0.2771 | 32.8/31.6 | -0.000 |
| 20170214 | L | -0.624 | 0.3961 [0.3261, 0.5014] | 0.2632 | 53.5/46.5 | -0.228 |
| 20170214 | Q | -328.107 | -52.7961 [-70.6504, -38.5864] | -50.0299 | 761.2/820.0 | -380.903 |
| 20170214 | B | -0.043 | 0.0841 [0.0648, 0.1103] | -0.0300 | 42.8/41.1 | 0.042 |
| 20170215 | L | 0.094 | 0.1052 [0.0875, 0.1238] | -0.0767 | 28.1/26.4 | 0.199 |
| 20170215 | Q | -0.366 | 0.1698 [0.1229, 0.2242] | -0.0636 | 34.5/32.3 | -0.197 |
| 20170215 | B | -0.366 | 0.1698 [0.1229, 0.2242] | -0.0636 | 34.5/32.3 | -0.197 |
| 20170216 | L | -1.907 | -0.0874 [-0.1833, -0.0194] | -0.2815 | 52.2/53.0 | -1.995 |
| 20170216 | Q | -915.677 | 6.5498 [3.9826, 10.2640] | 6.2908 | 927.0/923.7 | -909.127 |
| 20170216 | B | -0.694 | 0.0912 [0.0695, 0.1125] | -0.0167 | 39.8/38.8 | -0.602 |
| 20170217 | L | -62.967 | 16.9906 [11.4754, 25.7201] | 16.9163 | 147.6/126.5 | -45.976 |
| 20170217 | Q | -514481.0 | 17552.4 [11332.6, 27054.5] | 17352.3 | 13238.8/13011.0 | -496928.5 |
| 20170217 | B | -107.661 | 5.0822 [3.5329, 7.6641] | 2.0198 | 192.4/187.8 | -102.579 |
| 20170227 | L | 0.204 | 0.1065 [0.0848, 0.1258] | -0.1158 | 28.2/26.3 | 0.311 |
| 20170227 | Q | 0.187 | 0.0846 [0.0592, 0.1052] | -0.1281 | 28.5/27.0 | 0.272 |
| 20170227 | B | 0.194 | 0.0889 [0.0678, 0.1097] | -0.1250 | 28.4/26.8 | 0.283 |
| 20170228 | L | 0.201 | 0.1427 [0.1247, 0.1611] | -0.1182 | 37.9/34.4 | 0.344 |
| 20170228 | Q | 0.211 | 0.1279 [0.1117, 0.1469] | -0.1313 | 37.7/34.5 | 0.339 |
| 20170228 | B | 0.211 | 0.1279 [0.1117, 0.1468] | -0.1314 | 37.7/34.5 | 0.339 |
| 20170301 | L | -1.088 | 0.7599 [0.5802, 0.9920] | 0.4326 | 39.2/31.3 | -0.329 |
| 20170301 | Q | -11751.7 | 139.6246 [97.6998, 196.0392] | 152.9850 | 2940.0/2922.5 | -11612.1 |
| 20170301 | B | -7.919 | 2.8773 [2.1529, 3.7518] | 2.4062 | 81.0/66.7 | -5.042 |
| 20170302 | L | 0.123 | 0.1188 [0.1018, 0.1358] | -0.0671 | 27.0/25.1 | 0.242 |
| 20170302 | Q | 0.149 | 0.1020 [0.0872, 0.1168] | -0.0549 | 26.6/25.0 | 0.251 |
| 20170302 | B | 0.167 | 0.1014 [0.0866, 0.1159] | -0.0568 | 26.3/24.7 | 0.269 |

**Table S28.** Later Loco interval with position history. L, Q and B denote linear, quadratic and capped quadratic models. C/F gives baseline/full root mean squared error in mm/s. Gain intervals resample held-out blocks.

| Date | Basis | C $R^2$ | Gain [95% interval] | Replace | RMSE C/F | F $R^2$ |
|---|---|---|---|---|---|---|
| 20170210 | L | 0.258 | 0.0674 [0.0516, 0.0877] | 0.0320 | 23.9/22.8 | 0.325 |

| Date | Basis | C $R^2$ | Gain [95% interval] | Replace | RMSE C/F | F $R^2$ |
|---|---|---|---|---|---|---|
| 20170210 | Q | -453.080 | 158.4380 [112.5760, 218.3335] | 152.0255 | 590.5/476.4 | -294.642 |
| 20170210 | B | -0.859 | 0.2397 [0.1724, 0.3209] | 0.1731 | 37.8/35.3 | -0.619 |
| 20170213 | L | 0.272 | 0.0466 [0.0359, 0.0585] | -0.0204 | 27.0/26.1 | 0.318 |
| 20170213 | Q | 0.252 | 0.0423 [0.0275, 0.0574] | -0.0172 | 27.3/26.6 | 0.294 |
| 20170213 | B | 0.277 | 0.0443 [0.0299, 0.0588] | -0.0160 | 26.9/26.0 | 0.321 |
| 20170214 | L | -0.670 | 0.0664 [0.0450, 0.0964] | 0.0242 | 54.2/53.1 | -0.604 |
| 20170214 | Q | -1259.9 | -292.1766 [-386.5688, -218.5887] | -292.0261 | 1489.9/1653.6 | -1552.0 |
| 20170214 | B | -0.146 | -0.1010 [-0.1477, -0.0695] | -0.1182 | 44.9/46.9 | -0.247 |
| 20170215 | L | 0.554 | 0.0089 [0.0035, 0.0142] | -0.0193 | 19.7/19.5 | 0.563 |
| 20170215 | Q | 0.500 | 0.0227 [0.0143, 0.0341] | -0.0000 | 20.9/20.4 | 0.523 |
| 20170215 | B | 0.500 | 0.0227 [0.0143, 0.0341] | -0.0000 | 20.9/20.4 | 0.523 |
| 20170216 | L | 0.498 | 0.0342 [0.0258, 0.0457] | 0.0089 | 21.7/20.9 | 0.533 |
| 20170216 | Q | -197.447 | 108.3405 [66.4209, 164.2304] | 102.8513 | 431.3/290.6 | -89.107 |
| 20170216 | B | -1.133 | 0.1783 [0.1203, 0.2649] | 0.1552 | 44.7/42.8 | -0.955 |
| 20170217 | L | -88.043 | 33.0478 [22.2552, 50.3166] | 33.0535 | 174.2/138.1 | -54.995 |
| 20170217 | Q | -143243.9 | 10250.9 [6702.9, 15727.3] | 10198.1 | 6985.6/6731.0 | -132993.0 |
| 20170217 | B | -431.434 | 112.5300 [76.7606, 171.0161] | 112.4139 | 383.8/330.1 | -318.904 |
| 20170227 | L | 0.680 | 0.0112 [0.0074, 0.0150] | -0.0109 | 17.9/17.6 | 0.691 |
| 20170227 | Q | 0.684 | 0.0110 [0.0059, 0.0166] | -0.0083 | 17.8/17.5 | 0.695 |
| 20170227 | B | 0.689 | 0.0101 [0.0060, 0.0145] | -0.0080 | 17.6/17.3 | 0.699 |
| 20170228 | L | 0.666 | 0.0155 [0.0128, 0.0185] | -0.0118 | 24.5/23.9 | 0.681 |
| 20170228 | Q | 0.696 | 0.0123 [0.0096, 0.0151] | -0.0100 | 23.4/22.9 | 0.708 |
| 20170228 | B | 0.696 | 0.0124 [0.0097, 0.0151] | -0.0100 | 23.4/22.9 | 0.708 |
| 20170301 | L | -1.438 | 0.6650 [0.4690, 0.9174] | 0.6270 | 42.3/36.1 | -0.773 |
| 20170301 | Q | -4489.4 | 1118.4 [723.0561, 1631.3] | 1077.7 | 1817.3/1574.8 | -3371.0 |
| 20170301 | B | -5.138 | 1.3458 [0.9799, 1.7979] | 1.3117 | 67.2/59.4 | -3.792 |
| 20170302 | L | 0.635 | 0.0090 [0.0049, 0.0132] | -0.0090 | 17.4/17.2 | 0.644 |
| 20170302 | Q | 0.647 | 0.0097 [0.0062, 0.0136] | -0.0086 | 17.2/16.9 | 0.656 |
| 20170302 | B | 0.647 | 0.0097 [0.0063, 0.0136] | -0.0087 | 17.1/16.9 | 0.657 |

**Table S29.** Difference in later Loco mean squared error after capping elapsed time, in $(\mathrm{mm/s})^2$. Negative values indicate lower error for the capped model. T/H denotes task/position-history inputs.

| Date | Task context | Task full | History context | History full |
|---|---|---|---|---|
| 20170210 | -30368036.5 | -19781974.5 | -347208.1 | -225745.9 |
| 20170213 | -107.7556 | -113.2515 | -24.8366 | -26.8056 |
| 20170214 | -577606.0 | -670709.4 | -2217903.2 | -2732145.7 |
| 20170215 | 0.0000 | 0.0041 | 0.0000 | 0.0118 |
| 20170216 | -857758.8 | -851704.1 | -184036.3 | -82638.7 |
| 20170217 | -175229738.0 | -169251951.5 | -48651419.4 | -45197610.6 |
| 20170227 | -6.8333 | -11.1621 | -5.1200 | -4.3088 |
| 20170228 | 0.0000 | 0.0904 | 0.0000 | -0.0280 |
| 20170301 | -8637248.0 | -8536673.7 | -3298044.6 | -2476486.8 |
| 20170302 | -14.7325 | -14.2802 | -0.1788 | -0.1906 |
