## Supplementary material for "Motor cortical signals predict finger velocity beyond task geometry": Source data and analysis code: finger.pdf

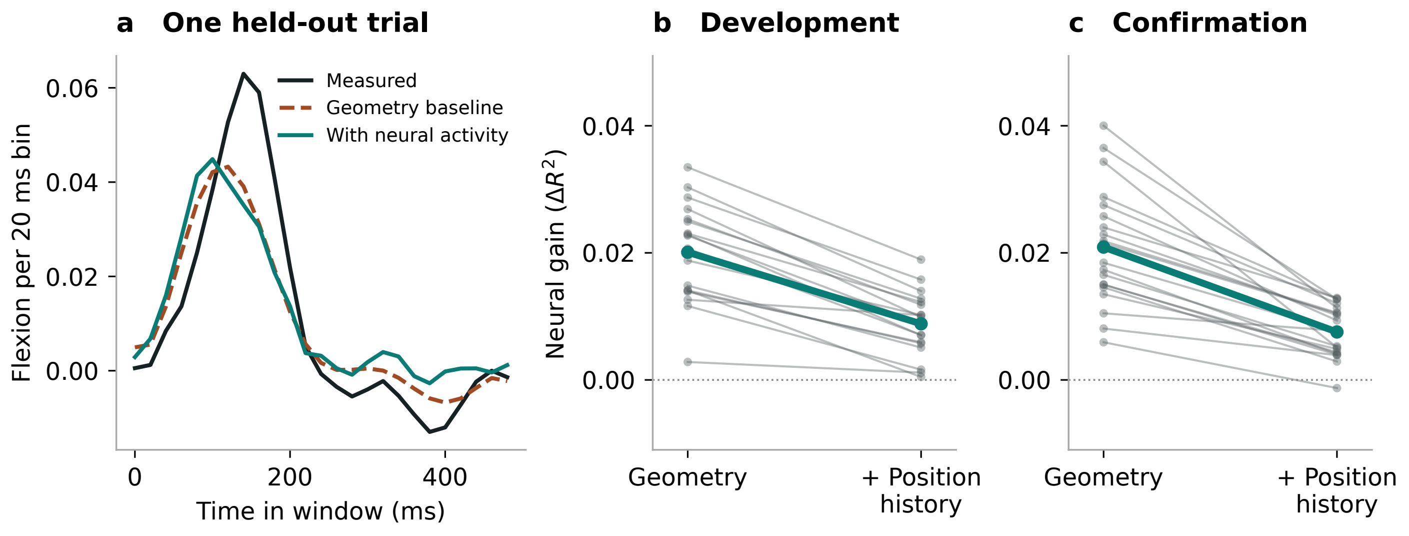

a One held-out trial
b Development
c Confirmation
Flexion per 20 ms bin
0.06
0.04
0.02
0.00
0
200
400
Time in window (ms)
Measured
Geometry baseline
With neural activity
Neural gain ($\Delta R^2$)
0.04
0.02
0.00
Geometry
+ Position history
