## Supplementary material for "Motor cortical signals predict finger velocity beyond task geometry": Source data and analysis code: pairing.pdf

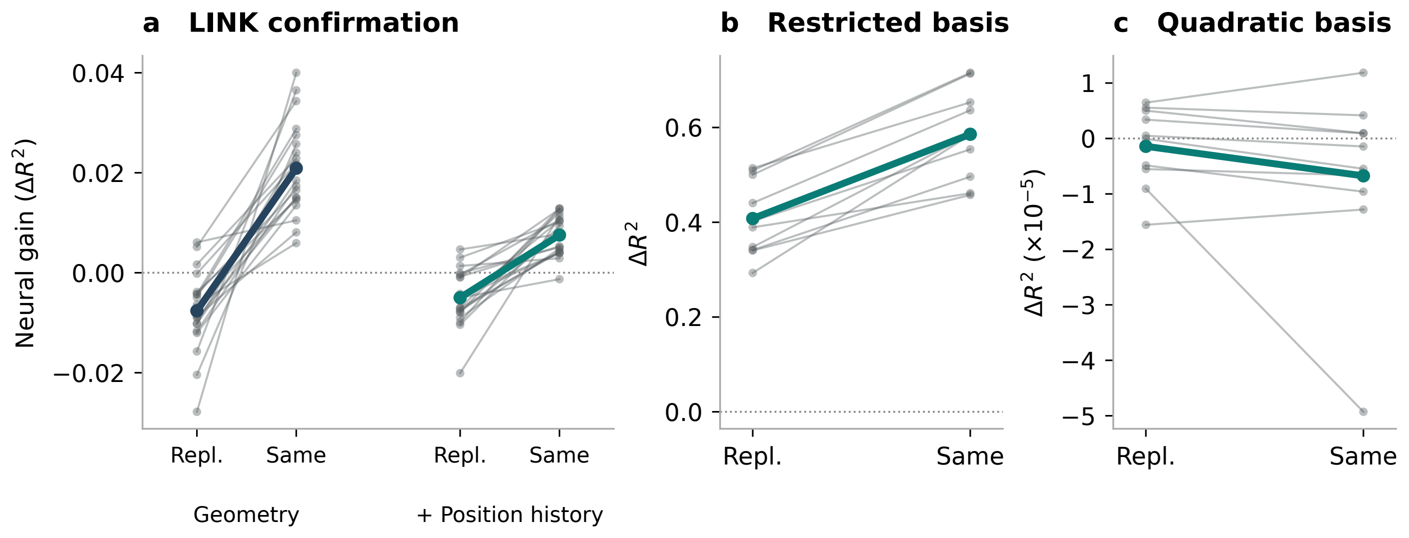
a LINK confirmation
Neural gain (ΔR²)
0.04
0.02
0.00
−0.02
Repl.
Same
Repl.
Same
Geometry
+ Position history
b Restricted basis
ΔR²
0.6
0.4
0.2
0.0
Repl.
Same
c Quadratic basis
ΔR² (×10⁻⁵)
1
0
−1
−2
−3
−4
−5
Repl.
Same
