## Supplementary material for "Motor cortical signals predict finger velocity beyond task geometry": Source data and analysis code: reaching.pdf

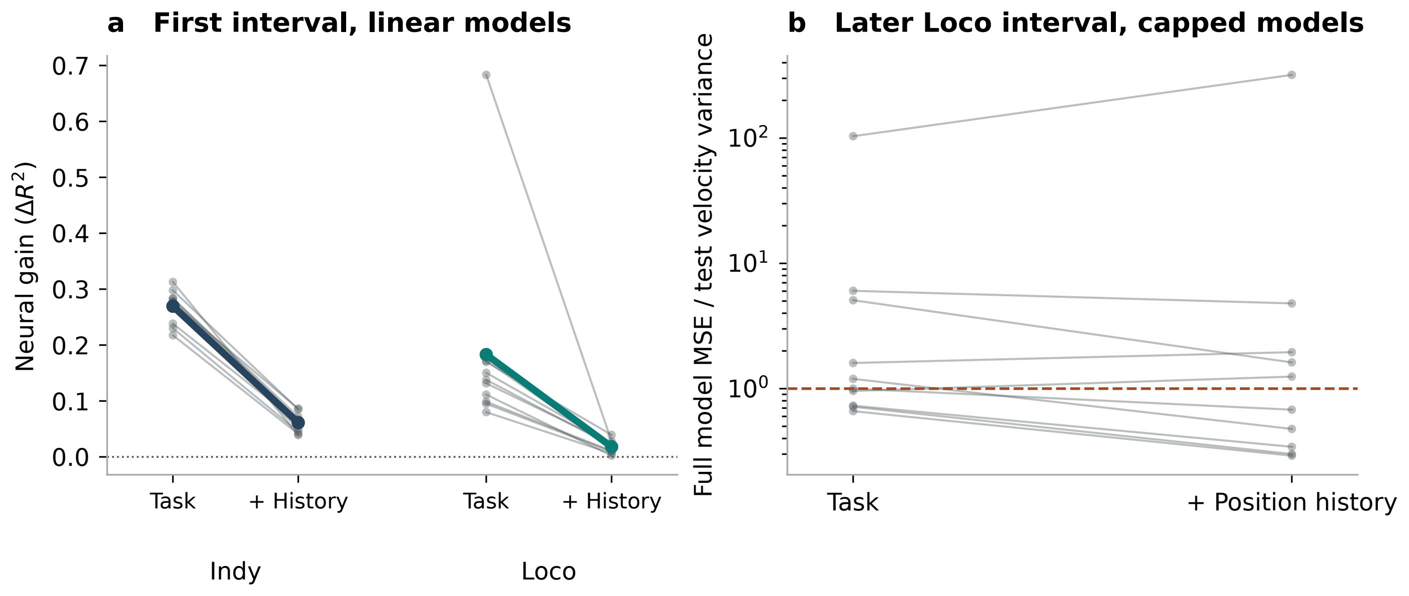

a First interval, linear models
Neural gain ($\Delta R^2$)
0.7
0.6
0.5
0.4
0.3
0.2
0.1
0.0
Task
+ History
Task
+ History
Indy
Loco
b Later Loco interval, capped models
Full model MSE / test velocity variance
$10^2$
$10^1$
$10^0$
Task
+ Position history
